# Prehistoric Global Migration of Vanishing Gut Microbes With Humans

**DOI:** 10.1101/2025.08.15.670570

**Authors:** Matthew M. Carter, Zhiru Liu, Matthew R. Olm, Melanie Martin, Daniel D. Sprockett, Benjamin C. Trumble, Hillard Kaplan, Jonathan Stieglitz, Daniel Eid Rodriguez, David A. Relman, Erica D. Sonnenburg, Michael Gurven, Benjamin H. Good, Justin L. Sonnenburg

**Author notes:** These authors contributed equally to this work (Matthew M. Carter, Zhiru Liu, Matthew R. Olm).

## Abstract

The gut microbiome is crucial for health and greatly affected by lifestyle. Many microbes common in non-industrialized populations are disappearing or extinct in industrialized populations. Understanding which microbes have been long-term residents of the human gut, and may have co-evolved with humans, has implications for the importance of microbial biodiversity loss for health. However, the genetic complexities of microbial evolution and the plasticity of gut microbiome composition have made it challenging to define these long-term associations. Here, we performed deep metagenomic sequencing of the Tsimane horticulturalists of Bolivia and compared their gut microbiomes with the Hadza hunter-gatherers of Tanzania. These two populations, whose ancestors were separated for tens of thousands of years, share 1,231 microbial species, most of which are absent in industrialized populations. Population genetic analyses in 636 of these shared species revealed patterns of microbial divergence and gene flow consistent with prehistoric human co-migration, with estimated split times that approximately align with human migration out of Africa and into the Americas. Our findings indicate that a diverse gut microbiome co-migrated with humans around the globe, persisting over millennia. However, many of these species are now vanishing in industrialized populations, and the consequences for human health remain uncertain.

## Introduction

Commensal gut bacteria play an intricate role in human biology^1^, but the evolutionary history of these microbial relationships remains poorly characterized^2^. Identifying long-term associations between humans and microbes is important for understanding their co-evolution^3–5^ and could reveal microbial species and functions integral to human health and physiology.

Insights into these microbial relationships have been gleaned from sequencing paleofeces^6^, with findings suggesting that ancient gut microbiomes were more similar to those of contemporary non-industrialized populations than industrialized populations (we use the term “industrialized” to refer to post-industrial populations as well as those in the ongoing and advanced stages of industrialization). However, the comparatively recent ages of existing paleofeces (∼1000-2000 years old), combined with the challenges of analyzing ancient DNA due to its degradation over time, makes it difficult to determine whether these microbes were also present in more ancestral human populations (∼10-100kya) or whether they reflect a more recent acquisition from a common environmental source.

Population genetic methods provide an alternative approach for inferring evolutionary history from the patterns of genetic diversity among contemporary microbial genomes^7–11^. Advances in strain-resolved metagenomics have made it possible to deploy this approach to gut microbial species sampled from human populations around the globe. Previous work has identified phylogenetic correlations between gut microbial genomes and their host populations of origin^5^, potentially consistent with a past history of co-migration. However, interpreting these correlations remains challenging due to the complex ways that microbial genomes evolve. Traditional phylogenetic methods^12^ break down for many species of gut bacteria, where horizontal gene transfer and homologous recombination produce mosaic genomes that cannot be represented by a single phylogenetic tree^13–16^. Moreover, existing strain-level analyses have been biased toward species in industrialized populations, which are rarely observed in ancient paleofeces. These sampling biases have hampered our ability to infer the evolutionary history of microbial taxa that are the strongest potential candidates for ancestral co-migration.

To address these challenges, we conducted deep metagenomic sequencing on fecal samples from the Tsimane horticulturalists of Bolivia (previously characterized with 16S rRNA sequencing^17^) and compared their microbiomes with those of the Hadza hunter-gatherers of Tanzania^18^. These two cohorts represent unique and geographically distant contemporary populations whose ancestors were separated thousands of years ago, providing a rare opportunity to examine human-microbe co-migration patterns across a large range of candidate ancestral microbial taxa. We identified over 1,200 microbial species shared between the two populations, most (60%) of which were absent in industrialized populations. Using population genetic analyses that explicitly account for microbial recombination, we provide evidence of strain-level reproductive isolation that is stronger than that observed in microbes from comparable industrialized populations. These findings suggest a transgenerational maintenance of gut microbial communities throughout ancient human migrations, offering new insights into the role of horizontal gene transfer and host-microbe co-evolution in the long-term structuring of the human microbiome.

## Results

### Metagenomic sequencing and genome recovery

The Tsimane are indigenous forager-horticulturalists living in the Bolivian Amazon. The individuals in this cohort live in six villages located along the Maniqui River and three additional villages in a nearby forest region^17^. We performed deep metagenomic sequencing on 133 previously collected fecal samples from 85 Tsimane adults sampled in 2009 and 2012-2013 (median depth = 31.9 gigabase pairs (Gbp); **Extended Data Fig. 1a**). We recovered 12,746 metagenome-assembled genomes (MAGs) representing 1,408 bacterial and archaeal species (**Fig. 1a, Methods, Supplementary Table 1**). We then compared these genomes to an analogous collection of 32,034 MAGs recently obtained from deeply sequenced microbiome samples of 137 Hadza hunter-gatherers in Tanzania^18^ (**Extended Data Fig. 1b**) to quantify how these populations’ gut microbial species have diverged over the last 100,000 years.

**Figure 1.**
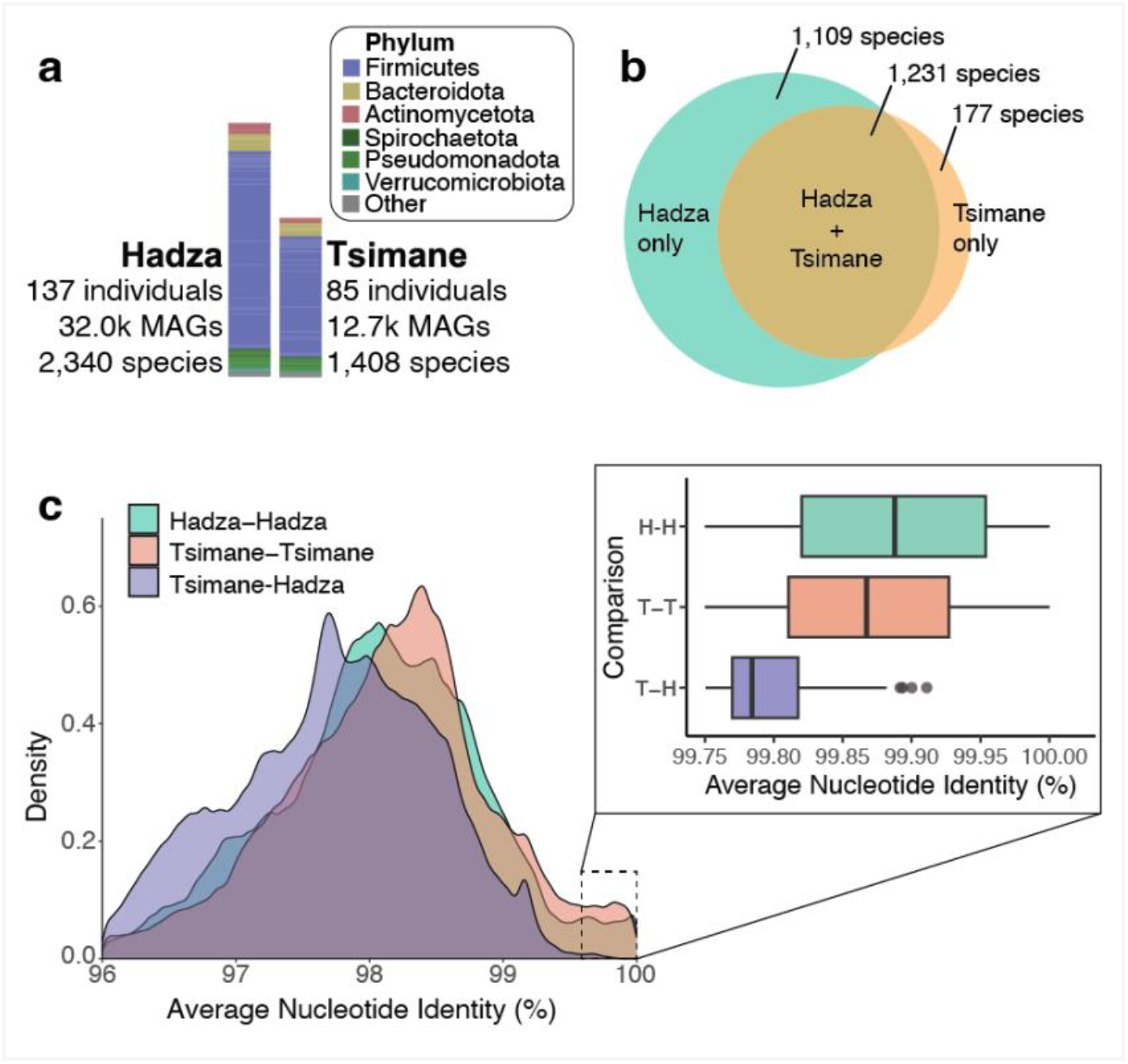
A large fraction of gut microbial species found in the Tsimane are also found in the Hadza. **a**, Stacked bar charts representing taxonomic diversity of microbial species recovered from Tsimane and Hadza microbiomes, colored according to bacterial phylum. **b**, Venn diagram depicting microbial species shared between populations. **c**, Density plot showing within- and between-population average nucleotide identity (ANI) across 541 species. Inset shows the corresponding distributions for ANI values greater than 99.75%.

### The Tsimane and Hadza share many species

Despite their large geographic separation and lack of historical direct contact, 87.4% (1,231) of the species in the Tsimane cohort were also found in Hadza individuals (**Fig. 1b**). Some of these shared species (e.g., *Agathobacter rectalis*) are also prevalent worldwide, but the majority (60.2%, 848) are undetected in the fecal microbiomes of industrialized populations (**Extended Data Fig. 2**). Species sharing between distant populations could arise through ancient co-migration, as well as more recent acquisition from the environment or human contact. We sought to distinguish these scenarios by examining the genetic variation within each of these shared species, and contrasting these patterns with the expectations from historical human migrations.

As a first step, we calculated the average nucleotide identity (ANI) between all pairs of MAGs within each of the 636 shared species with at least 4 MAGs in each population (**Extended Data Fig. 3**). These ANI values provide an upper bound on the time to the most recent common ancestor for each pair of MAGs. Estimates of the molecular clock in gut bacteria suggest that mutations should accumulate at a rate of at least ∼10^-7^/site/year (**Extended Data Fig. 4; Supplementary Methods**). The typical ANI between Tsimane and Hadza MAGs (∼98%; **Fig. 1c**) therefore requires that they must have shared a common ancestor within the last ∼100-200k years (**Extended Data Fig. 4**). The fact that most Tsimane species share a common ancestor with the Hadza on this timescale places strong constraints on their previous evolutionary history, and suggests that the ancestors of the Tsimane did not acquire a completely new suite of “local” gut microbes after their migration to the Americas.

Further insights can be obtained by comparing the ANI values between Tsimane-Hadza (T-H) pairs with their corresponding within-population counterparts (T-T, H-H). As expected, these data revealed that genomes from the same population tended to be more similar on average than genomes from different populations (P < 0.001, ANOVA, **Fig. 1c**). However, the effect sizes were small compared to the variation within each population (mean F_st_ = 0.10; **Supplementary Methods**). The broad overlap between these distributions indicates that most of the diversity within the Tsimane population is much older than the onset of geographic isolation.

### Reduced strain sharing in the recent past

While the average ANI values exhibited little geographic differentiation, we observed a much stronger trend in the tail of the distribution with high ANI (>99.75%), which shows a marked enrichment of pairs within versus between populations (P < 2.2 x 10^-16^, Wilcoxon rank-sum test, **Fig. 1c inset**). This within-population enrichment suggests that there was restricted strain transmission between these populations in recent evolutionary history, consistent with divergence driven by historical separation rather than ongoing exchange. However, estimating the timing of this separation can be challenging, since these high ANI values are often strongly influenced by recombination with other strains^13,16,19^. Previous work has shown that the high-ANI tail of distributions like **Fig. 1c** is composed of nearly clonal strains whose genomes have not yet been overwritten by recombination (**Extended Data Fig. 4; Supplementary Methods**)^13,16,20^. Their clonal ancestry can be inferred from their large stretches of nearly identical DNA sequences, which are interspersed with recombined segments from other strains. We estimated the total amount of clonal ancestry for each pair of MAGs using the fraction of core genes with identical DNA sequences (**Fig. 2a**). By calibrating these estimates with the rate of mutation accumulation in the vertically inherited regions of the genome, we determined that pairs of MAGs with >10% identical genes correspond to clonal strains that shared a recent common ancestor within the last ∼5,000 years (**Fig. 2a,b, Supplementary Table 2, Supplementary Methods**). Using this threshold, we identified all such pairs of clonal strains within the 636 shared species in **Fig. 1c** to investigate the landscape of recent strain transmission between the Tsimane and the Hadza.

**Figure 2.**
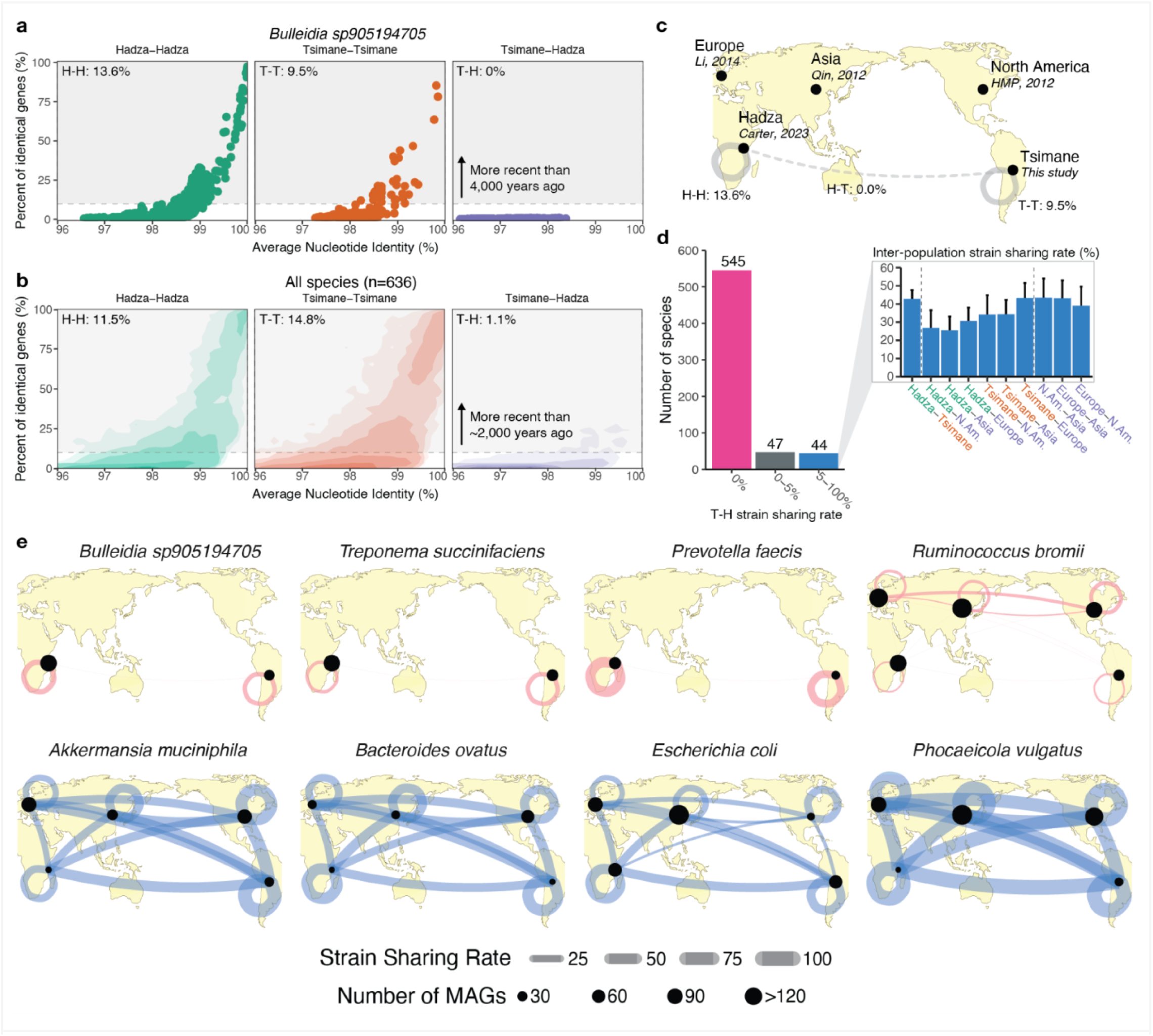
Most species shared between Tsimane and Hadza are genetically isolated, with the exception of some globally dispersed species. **(a, b)** Scatter plots showing the relationship between average nucleotide identity (ANI) and identical gene sharing between each pair of MAGs for (a) the species *Bulleidia sp905194705*, and **(b)** for 636 abundant bacterial species shared between the Tsimane and Hadza. Gray rectangles in each plot represent the 10% identical gene boundary used to define recently separated lineages; the age of this boundary varies across species, from ∼4,000 years for *Bulleidia sp905194705* to a median of ∼2,000 years (95% interpercentile range: 500-5000 years) for the larger group of species in panel **b (Fig. S3, S4 in Supplementary Methods)**. Percentages in each subplot indicate the percentage of MAG pairs that fall within the gray rectangle. **c**, Map showing location of the five cohorts we used for a global analysis of inter-population strain sharing. Gray lines represent the within (H-H and T-T) and between (T-H) population recent lineage sharing rate. **d**, Bar plot showing the number of species that have 0% Tsimane-Hadza (T-H) strain sharing rates, 0-5% T-H strain sharing rates and 5-100% T-H strain sharing rates. Inset shows the strain sharing rates for all inter-population comparisons for the 5-100% recent lineage sharing species. Error bars show standard deviation across species. **e**, Maps showing the within- and between-population sharing rates for eight species chosen to illustrate the range of different behaviors. The top four species have T-H strain sharing rates of 0%, the bottom four species have T-H strain sharing rates >5%. Size of dots represent the number of MAGs of that species derived from each population (no dot indicates no MAGs from that population). Thickness of connecting lines indicates the recent lineage sharing rate.

Aggregating comparisons across all 636 species revealed that, in total, 78,746 MAG pairs (7.8%) were clonally related (most recent common ancestor ≲5,000 years ago; **Figure 2b,c**). The vast majority of these clonally related pairs arose from intra-population comparisons (93.8%), indicating that most recent relationships occur within the same human population. Moreover, the relatively few clonal strain pairs shared between the Tsimane and Hadza were unevenly distributed across species, with 545 of the 636 (85.6%) species showing no recent T-H strain sharing (**Fig. 2d**). These results indicate that, in the vast majority of shared species, there has been very limited strain sharing between the Tsimane and Hadza in the past ∼5,000 years.

However, there were some notable exceptions to this trend of limited recent inter-population strain sharing. Some of the species exhibit levels of T-H strain sharing comparable to their within-population baselines. These species, such as *Akkermansia muciniphila* and *Bacteroides ovatus* (**Fig. 2e, bottom**), tend to be enriched in industrialized populations. To further explore this pattern, we expanded our analysis to include additional 8,855 MAGs from Europe, Asia, and North America^21–23^. The results revealed that these recently shared T-H species also had closely related MAGs distributed globally (**Fig. 2c-e, Extended Data Fig. 5**), suggesting that species with high T-H strain-sharing rates may have been introduced into the Tsimane and Hadza from more recent interactions with other human populations.

Intriguingly, the comparison with industrialized MAGs also revealed another class of species that are prevalent across lifestyles, but exhibit little recent strain sharing between the Tsimane and the Hadza (**Fig. 2e**). These taxa, which include common species like *Agathobacter rectalis* and *Ruminococcus bromii*, suggest that not all globally prevalent species are recent acquisitions from industrialized populations (or vice versa), and that global prevalence of a species does not always coincide with global strain transmission.

### Reduced gene flow via horizontal gene transfer

One limitation of strain sharing metrics is that the overall number of clonal pairs is shaped by multiple factors, such as host contact networks^24,25^ or genome-wide selective sweeps^26^, which can be difficult to compare across populations. To further corroborate our observation of microbial isolation between the Tsimane and Hadza **(Fig. 2)**, we also deployed an orthogonal approach that leverages information from the remaining pairs of MAGs (i.e. those with <10% identical genes). While their original clonal ancestry has been overwritten by recombination with other strains, recent recombination events between these more distantly related strains generate shorter segments of nearly identical DNA against a backdrop of lower genome-wide ANI (95-99%) **(Fig. 3a)**. Subsequent mutation and recombination events gradually erode these segments over time, creating a relationship between the length of an identical stretch of DNA and the age of the transfer event. The timing of these events can be leveraged to measure the rates of historical microbial gene flow within versus between human populations (**Fig. 3a**) – the classical genetic signature that they were part of the same microbial population^27^.

**Figure 3.**
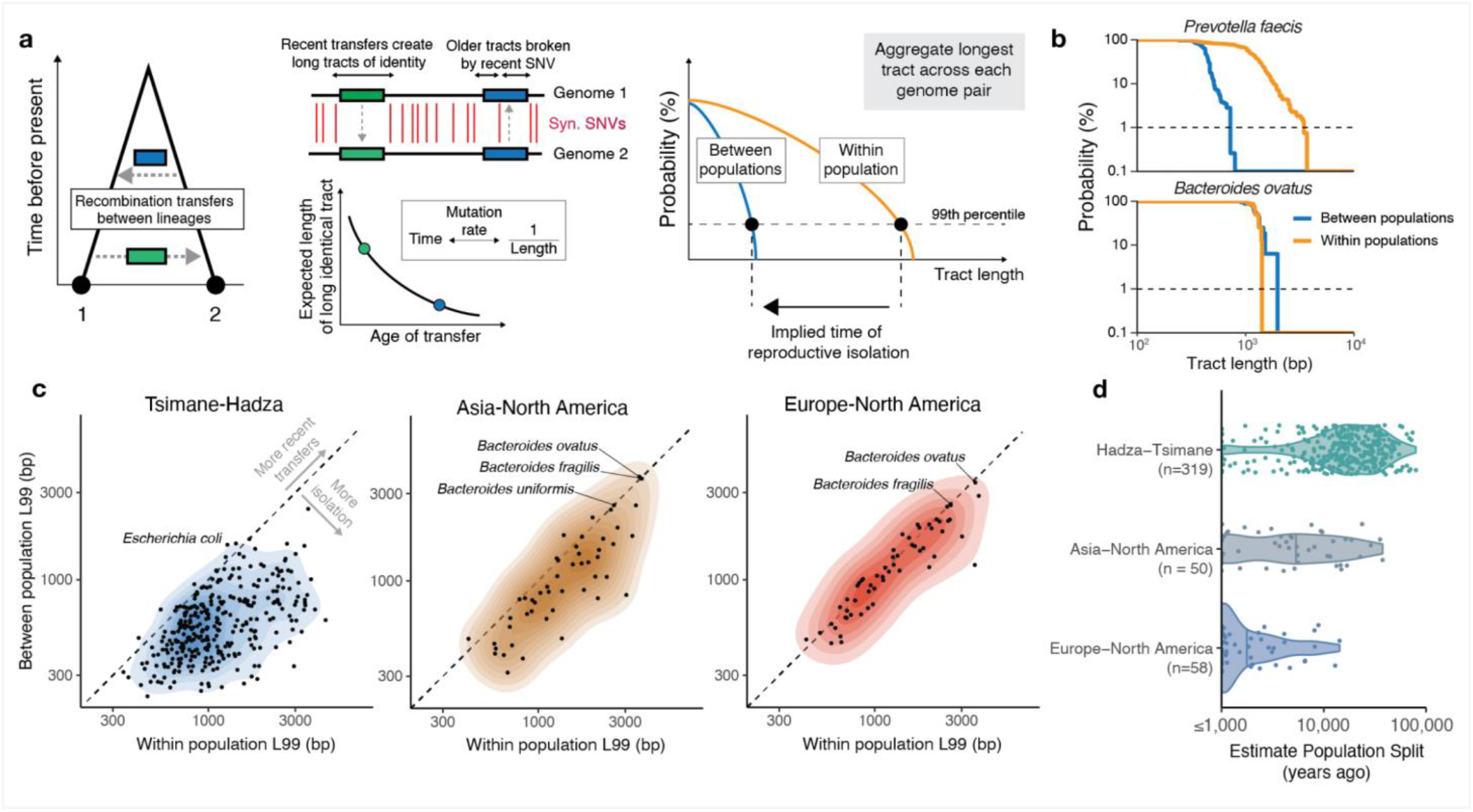
Lack of recent horizontal transfer of core genes is consistent with genetic isolation between Tsimane and Hadza gut bacterial populations greater than 10 kya. **a**, Recombination creates long identical genome tracts between strains, which are shortened by subsequent mutations, producing a characteristic relationship between tract length and transfer age (**Supplementary Note)**. Reproductive isolation causes long tracts between populations to derive from older, pre-isolation transfers, leading to shorter aggregated tract lengths compared to within-population comparisons. This difference can be used to infer the timing of genetic isolation. **b**, Empirical cumulative distribution function (ECDF) plots of tract length comparing two example species between and within the Tsimane and Hadza populations. Horizontal and dashed lines show the 99th percentile of the tract length distributions (L99 length). **c**, Comparison of the L99 metrics between several population pairs (Tsimane-Hadza, Asia-North America, North America-Europe). **d**, Violin plots showing the estimated population split time for each species, separated by the populations being compared.

We quantified this signal by identifying the longest tract of identical synonymous sites between each pair of MAGs within each of the 636 shared species above (representative species shown in **Fig. 3b)**. We excluded clonal pairs (>10% identical genes) to focus on gene flow rather than DNA acquired through a common ancestry (as analyzed in **Fig. 2**). Differences in tract lengths for within-versus between-population comparisons were most pronounced in the longer tracts, which coincide with recent recombination events (**Fig. 3b; Supplementary Methods**). Thus, to quantify this trend systematically, we computed the 99th percentile of tract lengths (“L99 length”) for each of the 636 species. Consistent with our previous results, most species had shorter L99 lengths between Tsimane and Hadza compared to within populations, suggesting genetic isolation in the recent past (**Fig. 3c**). However, exceptions such as *Escherichia coli* showed similar L99 lengths across populations, consistent with their high levels of more recent strain-sharing (**Fig. 2**). We repeated this analysis for MAGs from Asia, Europe, and North America (**Fig. 3c**). The differences in L99 lengths for within-versus between-population comparisons were smaller in these industrialized populations compared to the Tsimane and Hadza, suggesting a stronger barrier to gene flow between the Tsimane and Hadza compared to industrialized populations. This same trend was also observed within species that were prevalent across lifestyles (**Extended Data Fig. 6**), suggesting that it is not driven by systematic differences in the underlying microbial taxa being compared.

### Inferred microbial split times mirror ancient human migration timelines

Finally, we sought to use the observed L99 lengths in **Fig. 3c** to estimate the timing of the genetic isolation between the Tsimane and Hadza microbiomes. For each species, we added simulated mutations to pairs of within-population MAGs until their L99 lengths converged to their observed between-population levels. By calibrating these *in silico* population splits with the estimated mutation rates of gut commensals^16^, we can derive an approximate estimate for the age in years since a given pair of populations’ gut strains last exchanged DNA with one another **(Fig. 3d, Supplementary Table 3, Supplementary Methods)**. These estimates revealed that genetic exchange was notably more recent between industrialized populations (Europe-North America: median = 1,502 years, standard error (s.e.) = 732 years; Asia-North America: median = 8,430 years, s.e. = 2,378 years) than between non-industrialized populations (Tsimane-Hadza: median = 43,563, s.e. = 2,042 years). Of note, the calculated onset of genetic isolation between the Tsimane and Hadza microbiomes roughly aligns with the timeframe between the initial migration of humans out of Africa (approximately 60-70,000 years ago^28,29^) and the settlement of the Americas (approximately 16-23,000 years ago^30,31^), given the uncertainties in our molecular clock estimates (**Supplemental Methods**).

To obtain an additional independent estimate of microbial split times, we also employed an existing demographic inference method^32^. Rather than using stretches of identical DNA, this method uses an entirely orthogonal approach of analysing the distribution of single nucleotide variant (SNV) frequencies across populations (**Fig. 4a,b**; **Methods**). Although applying this model to microbes presents challenges, such as small sample sizes and complex population structures (e.g., the presence of clades or subspecies), we identified 15 species with sufficient data and appropriate population structure to estimate isolation times (**Supplementary Table 4, Supplementary Methods**). For each species, the inferred models broadly agree with those based on L99 lengths above **(Fig. 3d, Extended Data Fig. 7)** and clearly support an ancient population split in microbial demography rather than two populations with no isolation and free migration of microbes between populations (**Fig. 4b,c, Supplementary Methods**). Together, these results suggest that many gut microbial species have cohabitated with humans for at least as long as humans have been migrating around the world.

**Figure 4.**
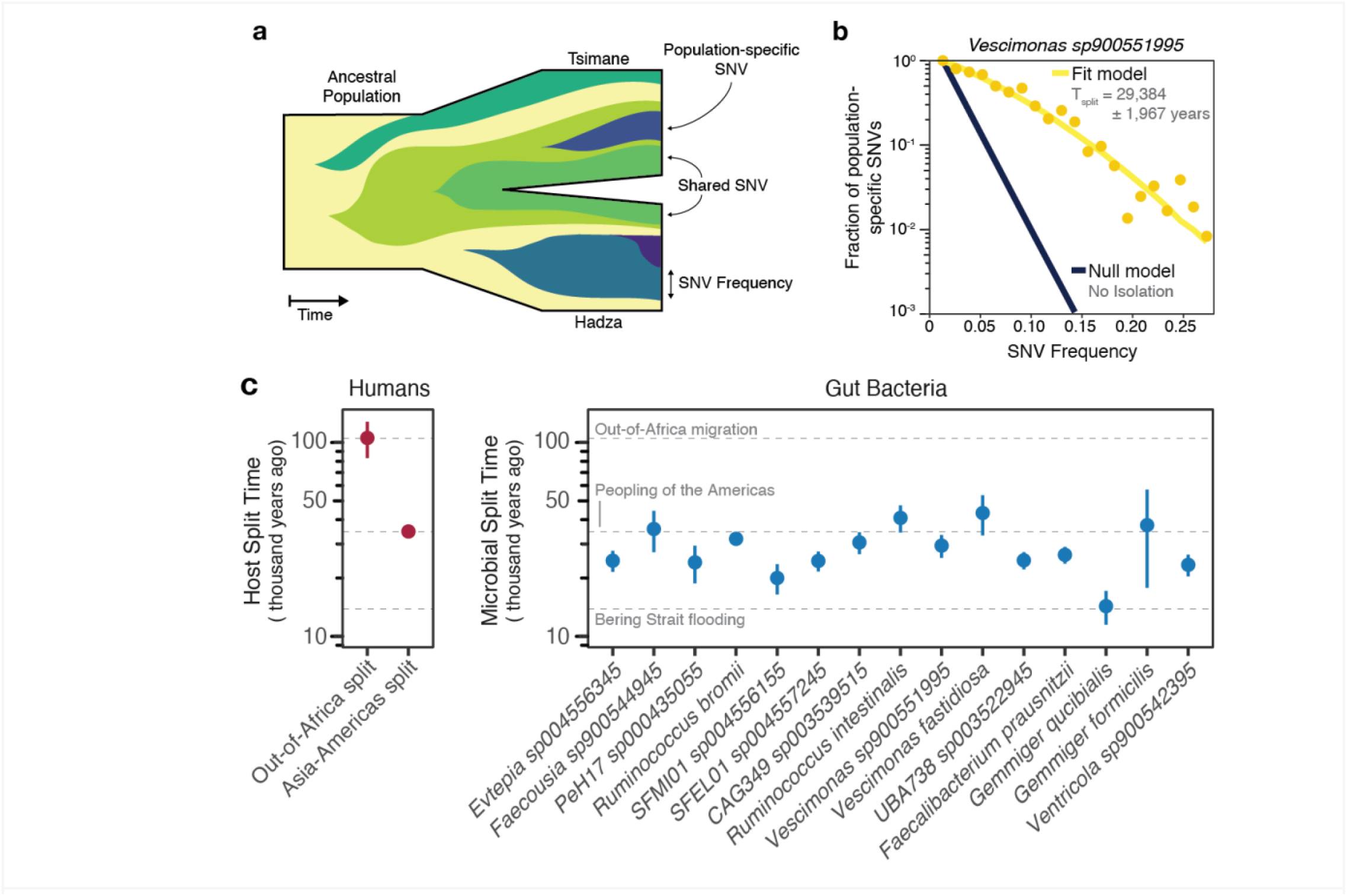
Gut bacteria show demographic patterns that mirror ancient human migration timelines. **a**, Schematic illustration of a population split, with each colored segment representing the allele frequency trajectory of a single nucleotide variant (SNV). More recent mutations are less likely to spread between genetically isolated populations. **b**, Comparison of demographic scenarios for one example species. The plot shows a slice of the full two-population SNV frequency distribution (**Supplementary Methods**), showing the relationship between SNV population frequency and the fraction of SNVs private to one population. Observed data from the Tsimane and Hadza cohorts (yellow dots) are compared with predictions from two scenarios: the inferred demographic model (yellow curve; uncertainty in T_split_ is the 95% confidence interval) derived using the full SNV frequency data and a no-separation scenario assuming free migration (black curve; **Supplementary Methods**). **c**, Inferred population split times of 15 gut bacterial species shared between the Tsimane and Hadza. For comparison, the inferred timings of two human population splits — the out-of-Africa migration and the Asia-Americas split — are shown on the left based on previously published work^33^ which used the same demographic modeling method^34^. The date of the Bering Strait flooding is based on Pico, et al.^35^ Error bars represent the estimated 95% confidence intervals from the model fit.

## Discussion

Using multiple, complementary analyses, we show that hundreds of commensal species inhabited the human gut for tens of thousands of years, transmitted over generations within Africa and during human migration around the world. These findings illuminate some of the historical dynamics shaping the present-day geographic distribution of gut microbiome species. Many of these species were previously identified as VANISH taxa (volatile and/or associated negatively with industrialized societies of humans) based on their enrichment in Hadza hunter-gatherers^18,36,37^. Our current genetic analyses demonstrate that many VANISH species were also present in the common ancestors of the Tsimane and Hadza, suggesting that widespread losses may explain these species’ absence from industrialized populations. Conversely, we also observed examples of BloSSUM species (bloom or selected in societies of urbanization/modernization)^38^ with extensive strain sharing across industrialized and non-industrialized populations, likely a result of contemporary species transmission from industrialized populations to the Tsimane and Hadza. Finally, we identified a new class of species (StIL; stable independent of lifestyle) prevalent across lifestyles, but exhibiting signatures of Tsimane-Hadza isolation that are consistent with prehistoric co-migration. Notably, within this subset of StIL species the degree of Tsimane-Hadza isolation was generally stronger than that observed between comparably separated industrialized populations (**Figure 2e, Extended Data Fig. 5**). These observations suggest that ongoing exchange within industrialized populations has likely eroded signals of co-migration, which may partially explain why previous observations of these signals have been limited.

These results highlight that, by focusing on the microbiomes of contemporary populations with limited exposure to industrialization such as the Tsimane and Hadza, we can better understand long-term human-microbiome associations that extend back to early human history. While these are contemporary populations, the genetic relationships between their gut bacteria can shed light on their past evolutionary history, similar to existing studies of the human genome^39^. As we expand our knowledge of microbiomes from rural and indigenous populations, it is critical to adhere to ethical research practices and foster collaboration with local communities, respecting their autonomy throughout the research process while ensuring mutual benefit, fostering open dialog, and building trust. Notably, the loss of VANISH taxa with changes in lifestyle, which are linked to an increased risk for a number of chronic metabolic and inflammatory disorders, raises the possibility that species reintroduction may help restore microbial functions that co-evolved with human biology and improve health^40^. This endeavor will require thoughtful, inclusive dialog among researchers, ethicists, and Indigenous partners.

## Acknowledgements

We are indebted to the participants in this study. We would like to thank the Hadza and Tsimane host villages and families that participated in this study as well as the numerous people and organizations who provided logistical support and conducted sample collection in Bolivia and Tanzania, namely the THLHP staff and researchers who provided invaluable assistance during field data collection, the Gran Consejo Tsimane, as well as Dorobo Safaris, John Changalucha, Alphaxard Manjurano, Maria Gloria Domiguez-Bello, Michelle St. Onge, Allison Weakley, Bryan Merrill, Samuel Smits, Yoshina Gautam, Dinesh Bhandari, Sarmila Tandukar, Guru Prasad Gautam, Jeevan B. Sherchand. The sequencing depth and breadth of this study was made possible by the Chan Zuckerberg Biohub, the Chan Zuckerberg Biohub Microbiome Initiative, and the UCSD Microbiome Sequencing Core. This work was funded in part by Open Philanthropy, a Stanford Bio-X Bowes Fellowship (to Z.L.), NIH/NIA R01-AG054442 (to H.K.), the Thomas C. and Joan M. Merigan Endowment at Stanford University (to D.A.R.), and NIH/NIGMS R35-GM146949 (to B.H.G.), J.S. acknowledges the French National Research Agency under the Investments for the Future (Investissements d’Avenir) programme (ANR-17-EURE-0010). J.L.S. and B.H.G. are Chan Zuckerberg Biohub Investigators.

## Author Contributions

Conceptualization: M.M.C., Z.L., M.R.O., B.H.G., and J.L.S. Methodology: M.M.C., Z.L., M.R.O., B.H.G., and J.L.S. Writing, review, and editing: M.M.C., Z.L., M.R.O., M.M.. D.D.S., B.C.T., H.K., J.S., D.E.R., D.A.R., E.D.S., M.G., B.H.G., and J.L.S. Visualization: M.M.C., Z.L., M.R.O., B.H.G., and J.L.S. Project administration: B.H.G., J.L.S. Supervision: B.H.G. and J.L.S. Funding acquisition: B.H.G., J.L.S.

## Data availability

Raw sequencing data generated as part of this study will be made available upon publication through the data access procedures established by the Tsimane Health and Life History Project (https://tsimane.anth.ucsb.edu/data.html). Additional datasets used in this study were downloaded from public repositories using the accessions provided in the cited references.

## Code availability

All custom code used for data analysis and figure generation is available at https://github.com/SonnenburgLab/comigration-metagenomics.

## Extended Data Figures

**Extended Data Figure 1.**
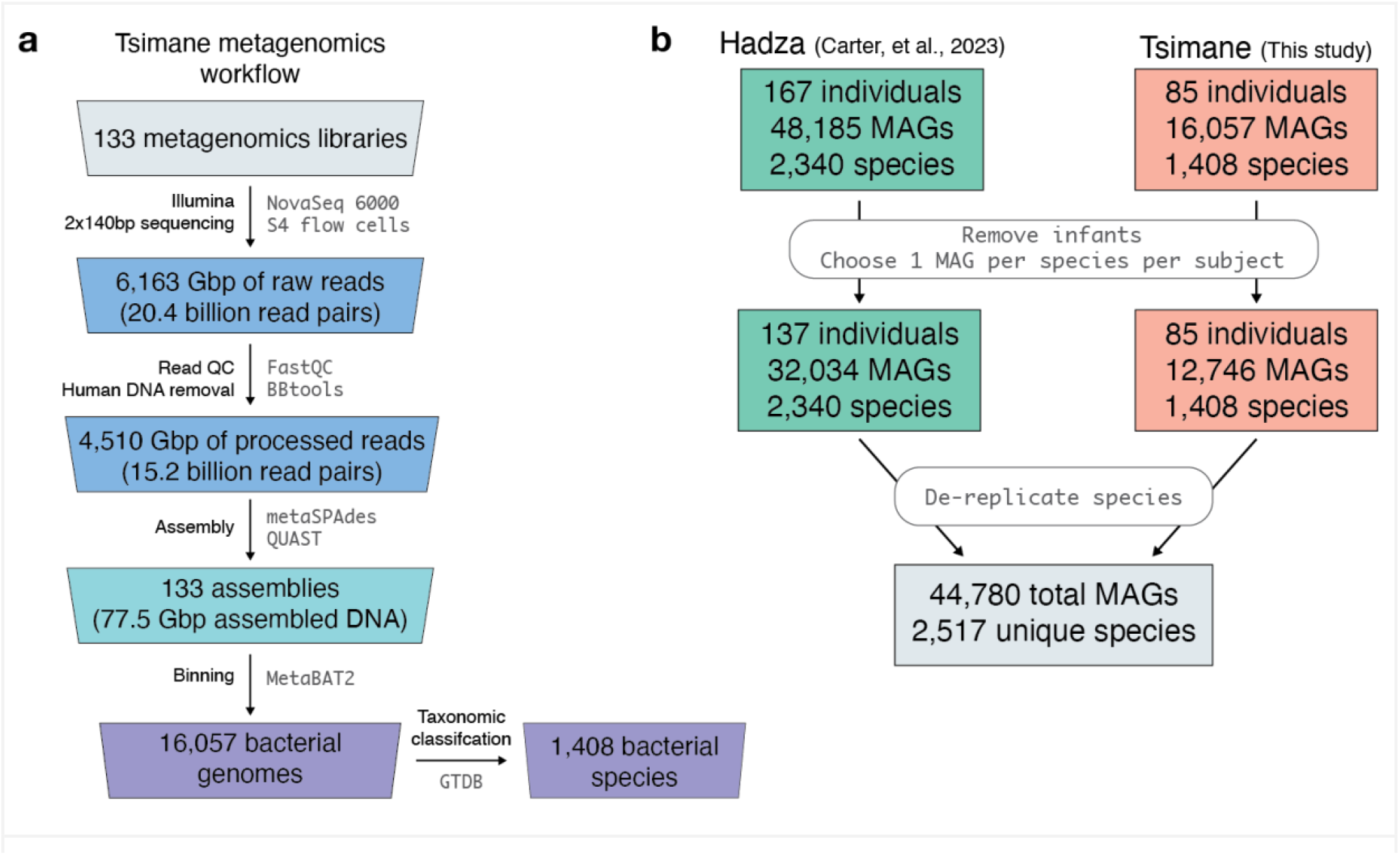
Metagenomics workflow and curation of Tsimane and Hadza data sets. **a**, Overview of computational workflow, tools used and primary data generated from Tsimane stool samples. **b**, Overview of genome database curation for Tsimane and Hadza cohorts.

**Extended Data Figure 2.**
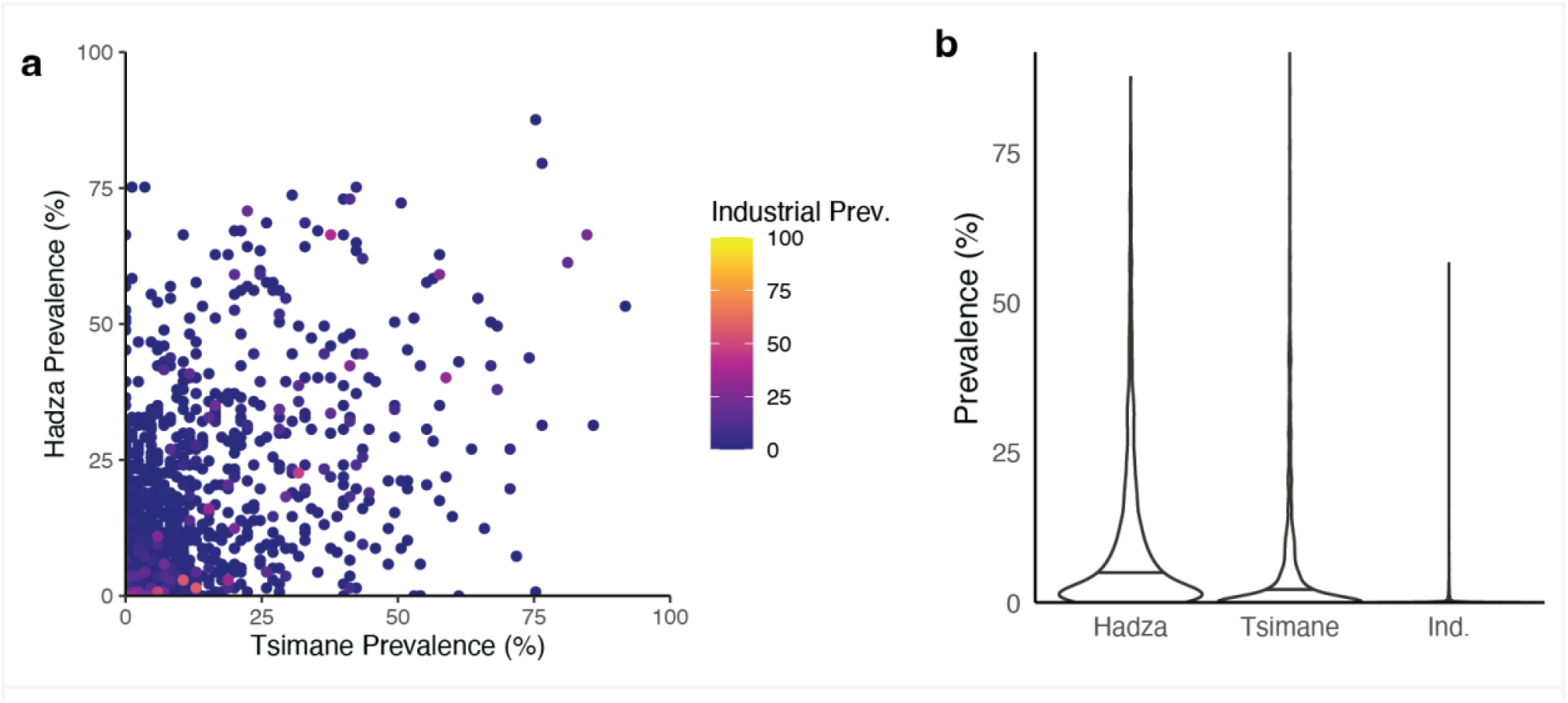
Prevalence of gut bacterial species in the Hadza, Tsimane and Industrial populations. **a**, Scatter plot showing the prevalence of the 1,440 species recovered in the Tsimane (x axis) and in the Hadza (y-axis) as well as industrial populations (color scale). **b**, Violin plots summarize the prevalence of gut microbial species in the Hadza, Tsimane and Industrial populations.

**Extended Data Figure 3.**
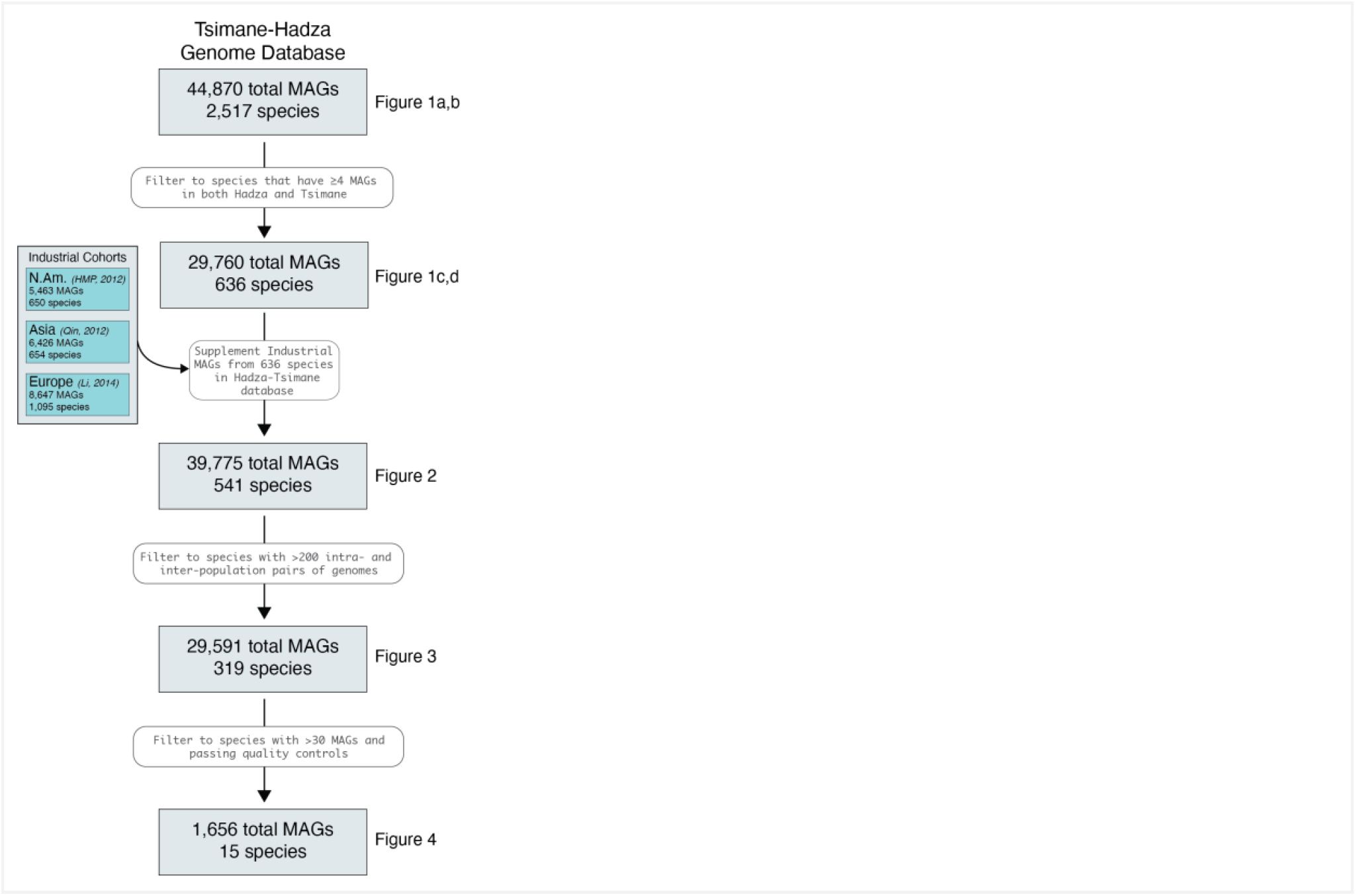
A schematic demonstrating which cohorts, MAGs and species were used for each analysis and the filtering criteria that were deployed. See **Supplemental Information** for quality control metrics for demographic inference.

**Extended Data Figure 4.**
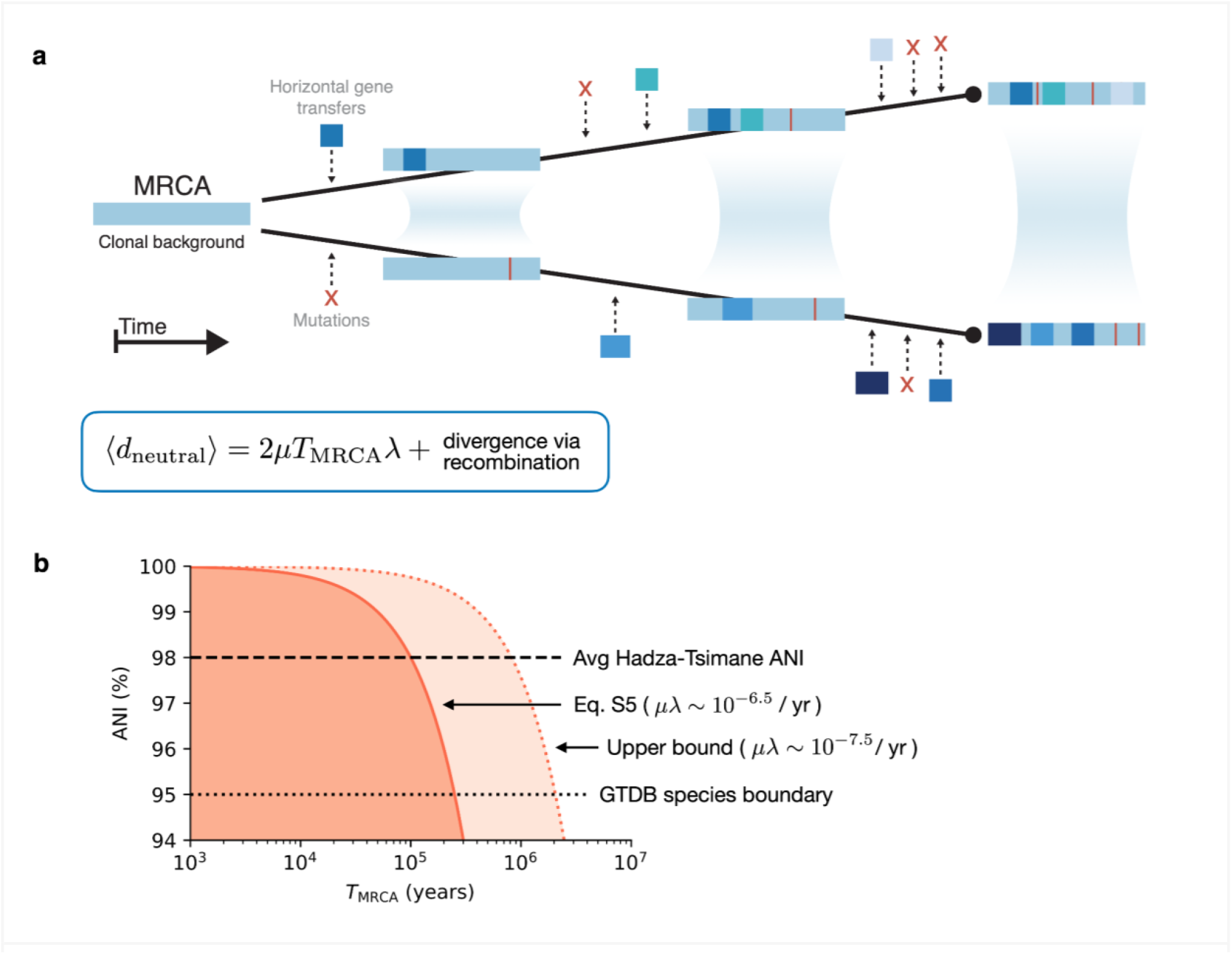
**a**, Schematic illustration of the bacterial molecular clock. A pair of strains descended from a clonal ancestor at time t=0 accumulate genetic differences via (i) de novo mutations (red) and (ii) homologous recombination events from other strains (dark blue). These differences accumulate as the divergence time increases, reducing the fraction of identical genes (light blue). **b**, Neglecting recombination leads to an upper bound on the expected time to the most recent common ancestor of a pair of MAGs (**Supplementary Methods**). Shaded regions denote the T_MRCA_ values that are consistent with a given ANI level. The solid red curve uses the midpoint estimate of the yearly mutation rate, while the dashed red curve denotes an approximate upper bound.

**Extended Data Figure 5.**
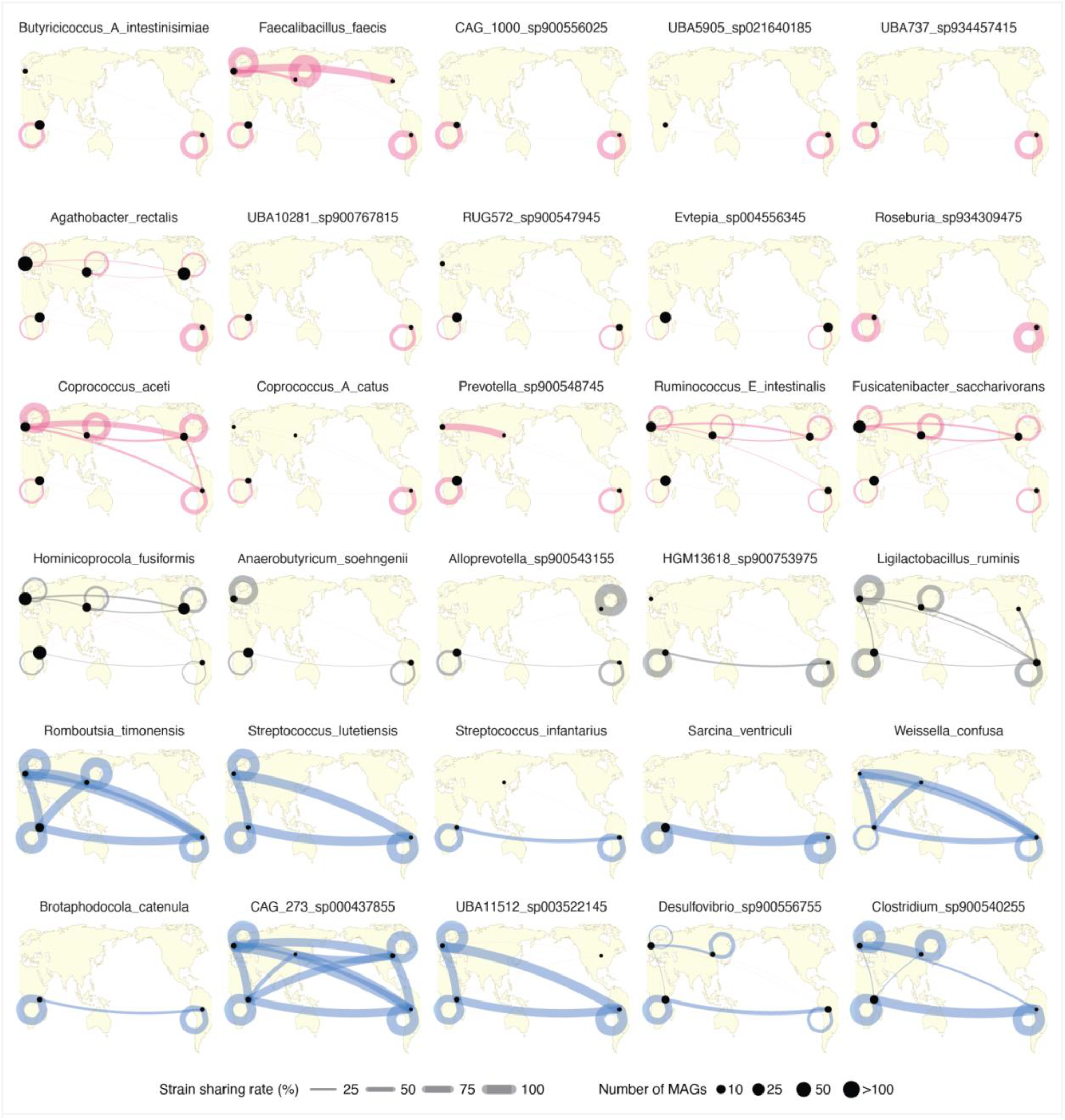
A larger collection of map plots, similar to the ones shown in Fig. 2e. The top three rows (magenta lines) have Tsimane-Hadza (T-H) strain sharing rates of 0%, the fourth row (gray lines) shows species with T-H strain sharing rates between 0% and 5%, the bottom two rows (blue lines) show species that have T-H strain sharing rates >5%. Size of dots represent the number of MAGs of that species derived from each population (no dot indicates no MAGs from that population). Thickness of connecting lines indicates the recent lineage sharing rate.

**Extended Data Figure 6.**
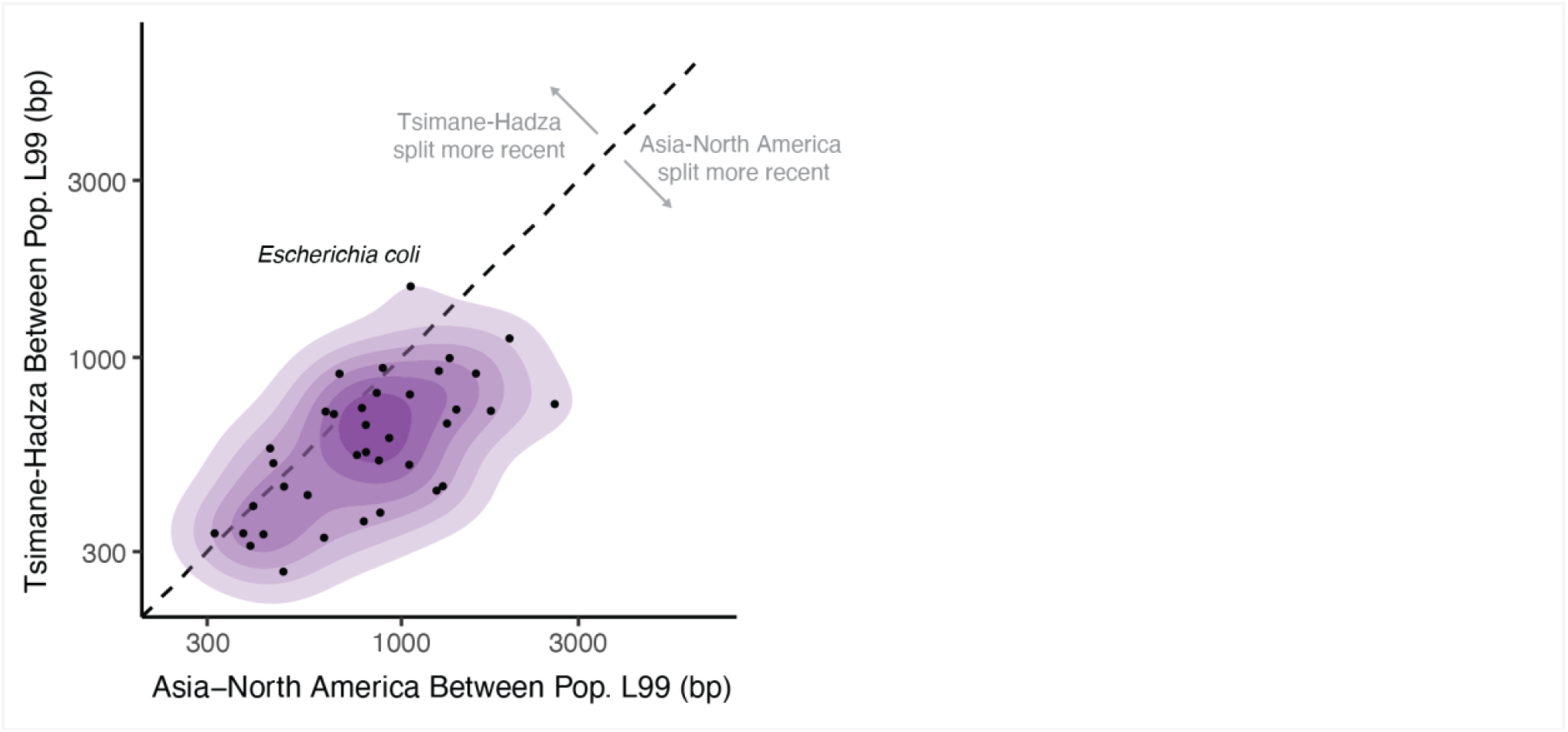
Comparison of the gut bacterial gene flow metric between the Tsimane and Hadza and between Asia and North America.

**Extended Data Figure 7.**
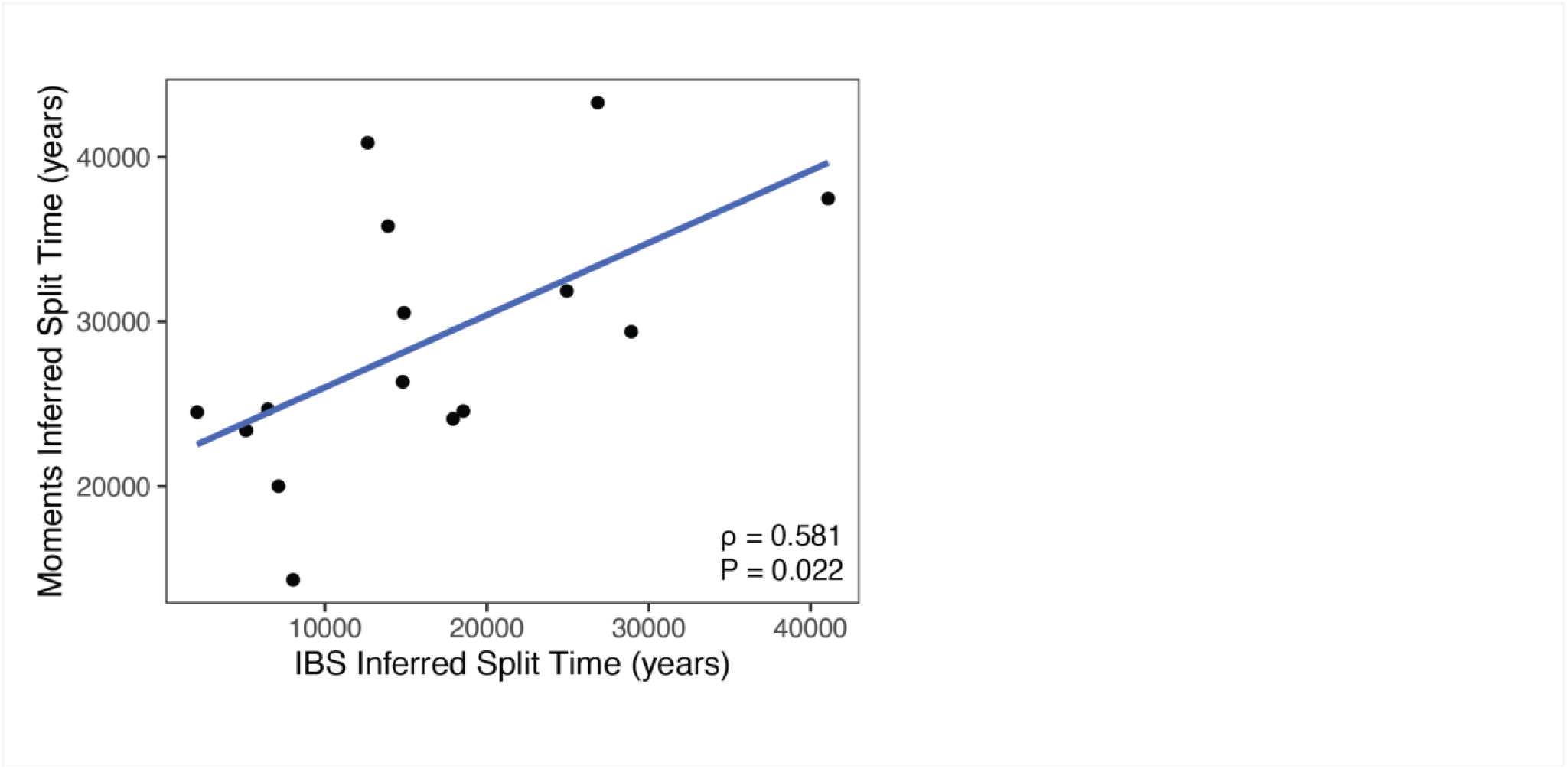
Scatter plot showing the correlation between the identity-by-state (IBS) inferred split time in years with the split time inferred by Moments demographic modeling for the 15 species for which we have data for both analyses (**Supplementary Methods**). Pearson correlation coefficient = 0.581, P = 0.022.

## Methods

### Ethics Approval

All study protocols were approved by the University of California, Santa Barbara Institutional Review Board on Human Subjects (IRB Protocols # ANTH-GU-MI-010-3U, submission ID 09-312, approved 8/21/2009; ANTH-GU-MI-010-19N, submission ID 12-354 approved 6/08/2012) and the Universidad Mayor San Simon, Cochabomba Bolivia. Permission to conduct research was granted to the Tsimane Health and Life History Project (THLHP) and their research affiliates. The THLHP maintains formal agreements with the local municipal government of San Borja and the Tsimane governing body. Consent was obtained from village leaders and community members during initial meetings upon starting research activities in each village. Consent was then obtained verbally from study participants prior to data collection.

### Sample collection

The Tsimane stool samples presented here were a part of previously published study^17^ and as part of the current work were subjected to additional sequencing. Briefly, the two sets of samples (2012–2013 and 2009) were collected across nine villages under the auspices of the THLHP^41^, which collects health and demographic data from Tsimane participants while providing primary health care to communities. The set of samples in this manuscript consists of Tsimane individuals at least 3 years of age (mean age = 28.4 years, s.d. = 13.2 years). Field specimen collection protocols were devised with the logistical challenges of this population in mind, and were consistent across the 2012–2013 and 2009 cohorts. At the time of data collection, none of the families in participating villages had plumbing, used pit toilets or diapers, or had access to refrigeration^17^. Fecal samples were collected in sterile urine specimen cups. Cups were given to individuals the day before sample collection. Researchers returned to participants’ homes between 7 a.m. and 9 a.m. to collect the specimens. Samples were transported to a field laboratory in coolers with reusable ice packs within 1–2 h of collection. Samples were homogenized in the collection cup and then partitioned into 2 ml sterile cryotubes using non-sterile wooden tongue depressors. Cryotubes were immediately stored in liquid nitrogen and then transferred to -20C freezers before transport to the U.S. on dry ice. Samples were stored in -80C laboratory freezers in the U.S. until analysis.

### Library preparation and sequencing

DNA was extracted from stool samples using MoBio PowerSoil kits (Qiagen, USA) following the manufacturer’s instructions. Libraries were prepared using Nextera Flex kits with a target of 10 ng of DNA with 12 base pair unique dual-indexed barcodes for 12 cycles to minimize amplification bias. Paired-end sequencing (2×140bp) was performed on a NovaSeq 6000 using S4 flow cells at the UCSD Microbiome & Metabolomics Core (San Diego, CA, USA). Samples were randomized across runs and sequenced repeatedly until the target depth was reached. Minimum target depth for each sample was 100 million paired-end reads (∼28 Gbp). A total of 5.25 terra base pairs of metagenomic data was generated.

### Metagenome quality-control and assembly

Raw sequencing reads were demultiplexed and processed using a custom processing pipeline (https://github.com/MrOlm/nf-genomeresolvedmetagenomics). Briefly, adapters were trimmed using FastP ^42^. Reads that mapped against human and PhiX genomes were removed using Bowtie2^43^. FastQC was used to ensure read quality^44^. Metagenomes were assembled using SPAdes^45^ and reads were mapped back against assemblies using Bowtie2. Contig coverage was determined using CoverM^46^. Assembly size and contig metrics were evaluated using QUAST ^47^ and assemblies were filtered to contigs >= 1500bp or all downstream analyses.

### Metagenome-assembled genome recovery

Genome binning was performed using METABAT2^48^ and genome bin quality was assessed using CheckM ^49^. Genomes with ≥50% completeness and <10% contamination according to CheckM were retained, in accordance with MIMAG standards^50^. Genomes with ≥50% completeness and <10% contamination were deemed “medium quality” genome bins and genomes with ≥90% completeness and <5% contamination were deemed “high quality” genome bins. We next used GTDB-Tk^51^ (r220) to determine the taxonomic classification of each medium- and high-quality genome bin. Genome bins derived from publicly available studies^21–23^ were also re-run through GTDB-Tk r220 to ensure up-to-date taxonomic classification.

### Strain divergence and recombination analyses

In order to determine the all-versus-all average nucleotide identity within each GTDB-assigned species, we used dRep^52,53^ v3.5.1 with the following parameters: (dRep compare --S_algorithm goANI --SkipMash). This command ensures that all genomes within a GTDB-assigned species are compared using the goANI algorithm^54,55^. Briefly, goANI identifies open reading frames using Prodigal v2.6.3^56^, aligns their nucleotide sequences with NSimScan^57^, and calculates ANI as the average sequence identity of all aligned genes.

To calculate the number of identical genes shared by a genome pair, the set of genes aligned between each genome pair was first filtered to include only those with at least 500 bp aligned. “Percent of identical genes” was then calculated as the number of gene pairs with 100% nucleotide identity divided by the total number of aligned gene pairs with at least 500 bp aligned. Further details on the methods used for the strain-sharing and recombination analyses in Figs. 2 and 3 are provided in the Supplementary Methods.

### Single nucleotide variant catalogs

For each species sufficiently represented in the Hadza and Tsimane we generated a single nucleotide variant (SNV) catalog of SNVs present in the core genomes of that species (core genomes are defined as the core genes present in >90% of MAGs within a species). To create an SNV catalog for a given species, we first chose one genome in the species as the representative genome based on which genome had the highest score according to the following formula: Completeness - 5*Contamination + 3*log10(N50) + Length/10^6^. We then used a previously published pipeline to annotate the sites in the representative genome that had SNVs compared to all of the other genomes in the species^58^ (https://github.com/zjshi/snv_analysis_almeida2019). We then filtered this whole genome SNV catalog to contain only sites that are in core genes. To determine which genes in each species’ pangenome qualified as a core gene, we called open reading frames on all genomes in the species using Prodigal v2.6.3^56^. We clustered all open reading frames in the species pangenome at 90% identity using MMseqs2 ^59^ and determined which gene clusters are present in at least 90% of genomes in the species. Based on these core genes, we created an additional mask that selected four-fold degenerate sites, thereby focusing on sites where all mutations would be synonymous. The resulting SNVs were used to perform the recombination and demographic inference analyses in Figs. 3 and 4. Further details on these analyses are provided in the Supplementary Methods.

