## Supplemental Information for "Prehistoric Global Migration of Vanishing Gut Microbes With Humans"

### Supplementary Information

**2 Assessing genetic differentiation with $F_{ST}$** **3**

**3 Estimating the divergence times of clonal strains** **4**

**4 Estimating population split times from recent HGT events** **5**
- 4.1 Identity-by-state (IBS) tracts . . . 5
- 4.2 Effects of population isolation . . . 6
- 4.3 Generating *in silico* population splits . . . 7
- 4.4 Application to empirical data . . . 7

**5 Demographic inference using Moments** **8**
- 5.1 Demographic model . . . 9
- 5.2 Quality control filters of model fits . . . 9
- 5.3 Visualizing model fits . . . 10

#### List of Figures

- S1 $F_{ST}$ between Tsimane and Hadza strains. . . . 14
- S2 2-D histogram showing the linear trend between the inferred clonal divergence time and the fraction of identical genes. . . . 15
- S3 Analogous versions of Fig. S2 for individual microbial species. . . . 15
- S4 Distribution of extrapolated clonal divergence times. . . . 16
- S5 IBS tracts of simulated microbial genomes using FastSimBac . . . 16
- S6 Validation of isolation-time inference using simulated data . . . 17
- S7 Comparison of within-population L99 scores for different populations . . . 18
- S8 Flowchart summarizing the filtering steps used for the Moments analysis . . . 19
- S9 Effects of clade structure in the Moments analysis . . . 20
- S10 Residuals from the Moments analysis . . . 21
- S11 Summary of Moments fits (1/2) . . . 22
- S12 Summary of Moments fits (2/2) . . . 24
- S13 Population-specific SNVs for simulated data . . . 25
- S14 Analogous versions of Fig. 4B for the remaining species. . . . 26

### Supplementary Methods

#### 1 Estimating the age of microbial isolation using the molecular clock

To convert the average nucleotide identity (ANI) measurements in Fig. 1 into a corresponding timescale, we employed a standard molecular clock approach informed by bacterial population genetics. We assume that the genetic differences in homologous regions of the genome emerge from a mixture of two processes: (i) point mutations, which alter individual sites; and (ii) recombination events, which replace longer stretches of DNA with a corresponding fragment sampled from another strain (Extended Data Fig. 4A). For a given pair of MAGs, the expected nucleotide divergence at neutral sites can then be expressed as a linear combination,

$$\langle d_{\mathrm{neutral}} \rangle = 2\mu \cdot T_{\mathrm{MRCA}} \lambda + \Delta_{\mathrm{r}}(T_{\mathrm{MRCA}}) \,, \tag{S1}$$

where $\mu$ is the spontaneous mutation rate (measured in units of mutations per bp per generation), $T_{\mathrm{MRCA}}$ is the time to the most recent common ancestor of the two strains (measured in years), $\lambda$ is the average growth rate (measured in units of generations per year), and $\Delta_r(T_{\mathrm{MRCA}})$ is a correction term that accounts for recombination events with other strains. While the precise form of $\Delta_r(T_{\mathrm{MRCA}})$ will depend on the nature of the underlying recombination events [1], one can show that under mild assumptions, this term is always greater than zero. Thus, neglecting recombination yields an upper bound on the expected time to the most recent common ancestor,

$$T_{\mathrm{MRCA}} \leq \frac{\langle d_{\mathrm{neutral}} \rangle}{2\mu\lambda} \,, \tag{S2}$$

which is illustrated in Extended Data Fig. 4B.

##### 1.1 Estimating neutral divergence from ANI measurements

We estimated the numerator of Eq. S2 under the standard assumption that most synonymous mutations are effectively neutral. However, since the ANI is an average over all sites, it is necessary to convert these measurements into a corresponding synonymous divergence rate. To do so, we decompose the ANI into a weighted average,

$$1 - \mathrm{ANI} = \frac{d_S \cdot S + d_N \cdot N}{S + N} = d_S \left( \frac{1 + \frac{N}{S}\frac{d_N}{d_S}}{1 + \frac{N}{S}} \right) \tag{S3}$$

where $d_S$ and $d_N$ represent the average divergence at synonymous and non-synonymous sites, respectively, and $S$ and $N$ are the corresponding numbers of synonymous and non-synonymous sites. While the $d_N/d_S$ ratio generally varies with ANI [2, 3], previous work has shown that the $d_N/d_S$ values for many species of gut bacteria are approximately ~0.1 when the ANI is $\lesssim$ 99% [3]. Similarly, assuming equal codon frequencies and an unbiased mutational spectrum, the ratio of non-synonymous to synonymous sites is approximately 3:1. Substituting these ratios into Eq. S3 yields an approximate conversion between ANI and the neutral divergence in Eq. S2,

$$1 - \mathrm{ANI} \approx d_S \left( \frac{1 + 3 \times 0.1}{1 + 3} \right) \approx \frac{d_{\mathrm{neutral}}}{3} \,, \tag{S4}$$

which applies for ANI values $\lesssim$99%.

#### 1.2 Estimating the yearly mutation rate

We estimated the denominator of Eq. S2 using two complementary approaches. *In vitro* measurements of bacterial mutation rates ($\mu$) range from $10^{-10}$ to $10^{-9}$ mutations per site per generation [4], while estimates of the growth rate ($\lambda$) in the human gut range from from 1-10 generations per day [5, 6]. Combining these estimates leads to a midpoint value of $\mu\lambda{\sim}10^{-6.5}$ per year, with a lower estimate of $\mu\lambda{\sim}10^{-7.5}$. More direct estimates have been obtained in two recent studies in *E. coli* [7] and *B. fragilis* [8], which measured the wall-clock substitution rate in longitudinally sampled isolates from individual human subjects. These studies yielded estimates of $\mu\lambda \approx 7\times10^{-7}$ and $\mu\lambda \approx 2\times10^{-7}$ per year for *E. coli* and *B. fragilis* respectively, which approximately coincide with the midpoint value ($\mu\lambda \sim 10^{-6.5}$ per year) from the first-principles calculation above. A recent study based on reconstructed genomes from ancient fecal samples likewise estimated $\mu\lambda \approx 4\times10^{-7}$ per year for *Methanobrevibacter smithii* [9]. In the absence of specific information for most of the taxa in Fig. 1, we adopted the mid-point estimate above for all of our subsequent analyses, with the understanding that the actual mutation rates may reasonably vary by ~2-3-fold in either direction.

#### 1.3 Application to the ANI measurements in Fig. 1

Substituting the above estimates into Eq. S2 yields a calibrated version,

$$T_{\mathrm{MRCA}} \lesssim \frac{100\times(1-\mathrm{ANI})}{2}\times10^{5}\ \text{years}\,, \tag{S5}$$

which is illustrated in Extended Data Fig. 4B. This approximate upper bound provides a benchmark for interpreting the ANI values in Fig. 1. Given the species definition employed by GTDB (ANI≥95%), two MAGs with the same species label must have shared a common ancestor within the last ~200-300k years. The average ANI between Tsimane and Hadza MAGs is even larger (≈98%, Fig. 1C), corresponding to a most recent common ancestor within the last ~100k years.

We note that the ANI estimates in Fig. 1 were only computed for MAGs in the same GTDB species bin; MAGs from non-shared species will have a more distant evolutionary relationship across populations. However, since more than 80% of the Tsimane species had a conspecific relative in the Hadza (Fig. 1B), we conclude that these ANI-based estimates must apply to the vast majority of the Tsimane gut microbiome.

We also note that the timescale in Eq. S5 should be interpreted cautiously. Given the uncertainties in the *in situ* mutation rates and ($\mu$) and growth rates ($\lambda$) in Eq. S2, the actual times may reasonably differ by several fold in either direction. However, these order-of-magnitude estimates still place strong constraints on the evolutionary history of the Tsimane strains. Taking the lower bound on the yearly mutation rate derived above ($\mu\lambda \sim 10^{-7.5}$ per year), we conclude that the Tsimane and Hadza MAGs must have shared a common ancestor within the last

$$T_{\mathrm{MRCA}} = \frac{3}{2}\cdot\frac{2\times10^{-2}}{10^{-7.5}} \approx 10^{6}\ \text{years}\,. \tag{S6}$$

This timescale is notably short compared to major geological events, or the origin of the *Homo* genus [10].

### 2 Assessing genetic differentiation with $F_{ST}$

To quantify the extent of population differentiation between the Tsimane and Hadza microbial populations, we applied the population genetic statistic $F_{ST}$, also known as the fixation index. $F_{ST}$ quantifies the proportion of total genetic variation that is attributable to differences between populations, as opposed to within-population variation. Values of $F_{ST}$ range from 0 to 1, where 0 indicates no differentiation (all genetic diversity is shared equally between populations), and 1 indicates complete differentiation (populations are entirely distinct). We estimated $F_{ST}$ using the formula:

$$F_{ST} \equiv \frac{d_{tot} - d_w}{d_{tot}} \equiv \frac{(1 - \mathrm{ANI}_{tot}) - (1 - \mathrm{ANI}_w)}{1 - \mathrm{ANI}_{tot}}\,, \tag{S7}$$

where $d_{tot}$ and $d_w$ represent the average pairwise genetic differences within the total population and within sub-populations, respectively. The observed $F_{ST}$ values typically ranges from 0.05 to 0.25, with a mean of 0.1 (Fig. S1). For comparison, the observed $F_{ST}$ between African and non-African human populations typically ranges from 0.1 to 0.3 [11]. This indicates that the differentiation observed between the microbial populations is relatively small.

In simplest model of a population split with no migration or external recombination, the expected value of $F_{ST}$ is given by

$$F_{ST} \approx \frac{T_{\text{split}}}{T_{\text{split}} + T_{\text{MRCA}}}\,, \tag{S8}$$

where $T_{\text{MRCA}}$ is the average time the most recent common ancestor for a pair of strains in the same population, and $T_{\text{split}}$ is the corresponding time since the two populations first diverged. While the actual mapping between $F_{ST}$ and $T_{\text{split}}$ is more complicated in more realistic demographic models (see Section 5.1 below), this result suggests that the genetic isolation of the Tsimane and Hadza microbial populations is relatively recent compared to the $T_{\text{MRCA}}$ of each species (100-200kya). More precise estimates of this split time are derived in Sections 3-5 below.

### 3 Estimating the divergence times of clonal strains

To quantify the timing of recent strain sharing between the Tsimane and the Hadza, we focused on pairs of clonal strains where the time of their most recent common ancestor can be inferred. Extended Data Fig. 4 shows that the genomes of these closely related strains will contain a mixture of clonal regions, which are nearly identical, and diverged segments imported from homologous recombination events with other strains. In the clonal regions, the recombination term in Eq. S2 vanishes,

$$\langle d_{\text{neutral}} \rangle \approx 2\mu \cdot T_{\text{MRCA}} \lambda\,, \tag{S9}$$

allowing us to directly estimate the divergence time of the strains using the molecular clock.

To distinguish the clonal regions of the genome from the recombined segments, we applied a recombination detection method (CP-HMM) from our previous study [1]. Briefly, this method models the spatial distribution of synonymous SNVs along the genome using a Hidden Markov Model. Because recombination introduces many nucleotide differences simultaneously, it results in localized clusters of SNVs. This elevated density of SNVs is the key signature exploited by the method to identify recombination events.

We applied this HMM-based method to 29,204 pairs of MAGs with more than 50% identical genes. Below this percentage, it becomes harder to resolve individual recombination events using our method because of a higher chance of overlapping recombination events. We then computed the average synonymous divergence within the clonal regions of each pair ($d_S^{\text{clonal}}$), and estimated the time to the most recent common ancestor using Eq. S9,

$$T_{\text{MRCA}} \approx \frac{d_S^{\text{clonal}}}{2\mu\lambda}\,, \tag{S10}$$

using the yearly mutation rate of $\mu\lambda \sim 10^{-6.5}$ mutations per site per year estimated in Section 1.2. The inferred $T_{\text{MRCA}}$ values for each analyzed pair shown in Figs. S2 and S3 and are provided in Supplementary Table 2.

While this method only directly infers the divergence times of strains with >50% identical genes, we extended this approach to more diverged clonal strains (10-50% identical genes) by extrapolating the observed relationship between divergence time and the fraction of identical genes. As a lower bound, we used the fact that the fraction of identical genes appears to decrease approximately linearly with clonal divergence for the data in Fig. S2. This linear relationship suggests a linear extrapolation to pairs of strains pairs with fewer than 50% identical genes:

$$f \approx 1 - cT_{\text{MRCA}}\,, \tag{S11}$$

where the constant of proportionality is equal to $c \approx 5 \times 10^{-4}$ year$^{-1}$. Under this model, the threshold we used for defining clonal strains – 10% identical genes – corresponds to an estimated divergence time of roughly 2,000 years.

As an upper bound, we used the fact that simple models of bacterial divergence [1] predict an exponentially declining relationship,

$$f \approx e^{-cT_{\text{MRCA}}}\,, \tag{S12}$$

which reduces to Eq. S11 when $1 - f \ll 1$. This exponential dependence accounts for the saturation that occurs when successive transfers start to overlap with each other. Fitting this model to the data yields an extrapolated divergence time that is ≈2.5x longer than Eq. S11 when $f \approx 0.1$. However, we note that the linear model appears to provide a better fit to the data even when $f \approx 0.5$, suggesting that the "true" extrapolation time may be closer to this lower bound.

We also performed the extrapolation separately for each microbial species (Fig. S3). We found that the estimated divergence time at a threshold of 10% identical genes varies between approximately 1,000 and 4,000 years under the linear model, with a median of about 2,000 years (Fig. S4). Together, these results support the conclusion that pairs of MAGs with >10% identical genes likely shared a common ancestor within the last ~2,000 years.

### 4 Estimating population split times from recent HGT events

#### 4.1 Identity-by-state (IBS) tracts

Identical segments between consecutive single nucleotide variants (SNVs) in a pair of genomes, also known as identity-by-state (IBS) tracts, contain valuable information about the demographic history of a population [12, 13]. IBS tracts arise from regions of recent common ancestry between the two genomes, either through direct vertical inheritance or recent recombination events between their ancestors. Regions with more recent ancestry tend to retain longer IBS tracts, as they have had less time for additional mutations and recombination events to accumulate. Consequently, the statistical distribution of IBS tract lengths reflects the underlying distribution of ancestry compositions, providing a means to infer past demographic processes such as population isolation [14].

Most previous studies of IBS tract length distributions have relied on models of crossover recombination in sexual organisms [15]. This make it hard to apply existing IBS methods to the microbial populations studied in this work, where recombination events involve shorter DNA segments [16]. In this section, we develop methods to quantify the signal of population isolation encoded in IBS tract lengths and estimate the isolation time through simulations.

#### 4.2 Effects of population isolation

Extended population isolation significantly affects the distribution of IBS tracts between populations. When a population splits into two isolated groups with no migration or gene flow, individuals sampled from different populations cannot share ancestry more recently than the population split time $T_{\text{split}}$. As a result, genome comparisons between the two populations will show fewer long IBS tracts compared to within-population comparisons, where genetic exchange continues. The magnitude of this reduction in IBS tract lengths provides quantitative insight into the duration of reproductive isolation between populations.

To focus on signals of recent demographic events, we recorded only the longest IBS tract between each pair of sequences. This approach improves computational efficiency and facilitates scaling to the full genome collection analyzed in this study. Throughout the remainder of this section, we use "IBS tract distribution" to refer to the distribution of these longest IBS tracts across genome pairs. We also excluded clonal pairs from this analysis (Section 3), since their longest IBS tracts are generated by vertical inheritance, rather than recent horizontal gene transfer. These latter events provide a more direct signal that two genomes are part of the same population unit, which is the goal of the present section.

To illustrate effect of population isolation on the IBS tract distribution, we utilized a population genetic simulator FastSimBac to simulate a simple neutral scenario of population split followed by complete isolation. FastSimBac generates genome sequences similar to our observed data, allowing us to compute IBS distributions within and between populations using the same procedures that we applied to the observed data. We selected simulation parameters designed to match the observed data of typical human gut commensals [1]. We set the genome length to $10^6$ basepairs, the population-scaled mutation rate to $\theta = 0.01$, the population-scaled recombination rate to $R = 0.005$, and the typical recombination length to $\ell = 10^4$. The simulated demographic scenario assumes that an initially well-mixed population of size $N$ splits into two populations ($N$ each) $T_{\text{split}}$ generations ago, after which they evolve in isolation. We simulated a sample of 50 individuals from each subpopulation across a range of scaled split times, defined as $\tilde{T}_{\text{split}} \equiv T_{\text{split}}\lambda/2N$, where $\lambda$ is the average growth rate. The exact simulation commands can be found in the associated analysis source code.

Figure S5 shows that, compared to within-population comparisons, IBS distributions between populations exhibit a sharp cutoff at a characteristic length scale, similar to the pattern in observed data (e.g. Fig. 3B in the main text). As isolation time increases, this characteristic length scale decreases, reflecting the accumulation of mutations and recombination events that breaks apart long IBS tracts.

To systematically quantify the signal of population isolation, we focused on the 99th percentile of the IBS tract distribution (the L99 length). Since the cutoff in the tail of the IBS distribution is sharp in both simulations and observed data, this statistic is robust to the exact choice of percentile. The right panel of Fig. S5 shows how the between-population L99 length decreases as a function of isolation time in FastSimBac simulations. The clear decay of the L99 length as $T_{\text{split}}$ increases suggests that we can infer the isolation time from the observed L99 lengths by inverting this functional relationship.

However, the precise form of the L99-$T_{\text{split}}$ curve will vary across species due to differences in biological parameters such as recombination rate and tract length. Additionally, we expect that real populations may have more complex demographic histories than the simple scenario modeled above (e.g., a population bottleneck or expansion after the split). Generating species-specific L99-$T_{\text{split}}$ curves through tailored FastSimBac simulations is therefore not only computationally demanding but also requires additional demographic assumptions for each species. Instead, we sought to derive these curves directly from observed genome sequences using a semi-parametric approach, which approximates the process of mutation accumulation after the population split.

#### 4.3 Generating *in silico* population splits

Our approximate simulation method leverages the observed genome sequences to model the effects of a population split, by simulating the accumulation of additional mutations over time. This semi-parametric approach avoids the need for computationally intensive, species-specific recombination simulations while incorporating species-specific parameters in a natural way. This method relies on two key assumptions. First, the within-population IBS distribution observed in present-day genomes must be a good approximation of the ancestral IBS distribution prior to the population split. Second, spontaneous mutations must be the primary factor breaking apart long IBS tracts over time, with recombination playing a secondary role over the length- and time-scales of interest.

For a given species, we estimated the L99 length versus isolation time relationship using the following procedure:

1. For each pair of sequences from the same population, we recorded the observed locations of synonymous SNVs and the total (core) genome length $L$.

2. For a given isolation time $T_{\mathrm{split}}$, we introduced $k$ new SNVs randomly along the genome, where $k$ is a Poisson random variable with mean $2\mu \cdot T_{\mathrm{split}}\lambda \cdot L$, with $\mu$ and $\lambda$ denoting the average mutation rate and growth rate from Eq. S2. We used the yearly mutation rate of $\mu\lambda \sim 10^{-6.5}$ mutations per site per year estimated Section 1.2.

3. We then computed the IBS tract lengths based on the new set of SNVs and recorded the longest tract for each pair.

4. We then repeated steps 1-3 for all within-population sequence pairs and a range of isolation times $T_{\mathrm{split}}$.

5. We then downsampled the data to match the observed number of between-population pairs for the species of interest and computed the L99 length for each simulated isolation time $T_{\mathrm{split}}$.

To test the accuracy of this approximation, we applied it to the FastSimBac simulation data above. We generated approximate L99-$T_{\mathrm{split}}$ curves using within-population pairs from a single simulation and compared it to the ground truth computed in Fig. S5. Figure S6 shows that our approximate simulation method accurately recapitulates the ground truth relationship across a range of isolation times. The estimated curve closely follows the ground truth, with minor underestimation of the L99 length likely due to unmodeled recombination effects. This agreement suggests that mutation accumulation alone provides a strong first-order approximation of IBS tract dynamics in our regime of interest, supporting the validity of this approach for estimating isolation times in empirical data.

#### 4.4 Application to empirical data

We used this approach to calculate IBS tracts and L99 lengths for all of the species in Fig. 3. To minimize the impact of purifying selection, we focused on fourfold synonymous sites within the core genes of each species (i.e. those present in >90% of MAGs, see Methods). To ensure that our analysis is independent of the clonal pair analysis in Fig. 2 (Section 3), we excluded genome pairs with >10% identical genes. For the remaining genome pairs, we concatenated contigs of the reference MAGs and recorded the spacings between consecutive SNVs. We recorded IBS tracts distributions for all pairs of MAGs within each population as well as between populations (Tsimane-Hadza, North America-Europe, North America-East Asia). We found that within-population L99 lengths are highly consistent across populations (Fig. S7), suggesting that this statistic is robust to sampling differences across studies. Consequently, we utilized the average within-population L99 length regardless of the specific population pair.

**Inferring the population split time.** We generated species-specific L99-$T_{\text{split}}$ curves by performing mutation accumulation simulations using the algorithm described in Section 4.3. We pooled all within-population genome pairs across all populations and excluded species with fewer than 200 such pairs. We simulated 20 different values of $T_{\text{split}}$ that were evenly spaced on a log scale between $10^3$ and $10^{5.5}$ years. For each population pair, we downsampled the simulated data to match the number of observed between-population genome pairs and computed the corresponding L99 length for the simulated value of $T_{\text{split}}$.

In practice, we found that at shorter isolation times, the simulated L99 lengths were often indistinguishable from the within-population L99 lengths due to sampling error. This observation suggests a lower bound of approximately $1,000$ years on the resolvable isolation time between populations.

To obtain a continuous function relating the L99 length to $T_{\text{split}}$, we applied isotonic regression to enforce monotonicity in the simulated results. This step, which was implemented using the Python package scikit-learn [17], reduces noise-related fluctuations, particularly for small values of $T_{\text{split}}$ where the L99 lengths are similar. Using this approach, we extracted the smoothed L99 lengths at the simulated values of $T_{\text{split}}$ and performed linear interpolation between $\log(L99)$ and $\log(T_{\text{split}})$. We chose to interpolate in log space to account for the power-law scaling observed in simulations (Fig. S6).

We inferred the implied isolation time between populations by solving for the point where the interpolated L99-$T_{\text{split}}$ curve intersected the observed between-population L99 length, using the default root-finding algorithm in scipy [18]. When the observed L99 length fell outside of the simulated range, we assigned the inferred isolation time as either below $10^3$ years or above $10^{5.5}$ years. In practice, no comparisons exceeded the upper range, while Europe-North America pairs frequently fell below the lower bound, suggesting no detectable isolation. This result is consistent with the near-identical within- and between-population L99 values reported in Fig. 3C of the main text.

To quantify the uncertainty in the inferred isolation times, we performed 1,000 bootstrap replicates for each population pair and calculated 95% confidence intervals. This resampling approach also allowed us to assess the statistical significance of population isolation. Specifically, if more than the 95% of bootstrap replicates produced an L99 value below the simulated range, we concluded that the species exhibited no detectable population isolation between the two populations. The final inference results are provided in Supplementary Table 3.

#### 5 Demographic inference using Moments

To corroborate the estimates above, we also employed an existing demographic inference method (Moments; 19) which leverages information from the joint site frequency spectrum (SFS) of SNV frequencies within populations. Moments is traditionally applied to diploid organisms, so we implemented several additional filtering procedures to adapt it to the microbial populations analyzed here.

To ensure sufficient resolution in the observed SNV frequencies for demographic inference, we focused on species with the largest number of MAGs. Specifically, we retained species with more than 30 MAGs in both the Tsimane and Hadza populations. For context, Medina-Muñoz et al. used 40 haploid copies per population in their inference of human demographic history [20]. We computed the site frequency spectrum for these selected Tsimane and Hadza MAGs using four-fold degenerate synonymous sites. As part of this process, we applied the projection function in Moments that standardizes total allele counts across sites by downsampling to a specified threshold. We set this projection threshold to 0.9 times the number of MAGs in each population, rounded down to the nearest integer.

Since bacterial MAGs derived from metagenomes may have higher rates of SNV calling errors compared to high-quality haplotypes from whole-genome sequencing (WGS) of human genomes (e.g. in Ref. 20), we sought to minimize their impact by masking singleton SNVs in each population during model fitting. Specifically, this was implemented by setting `data.mask[1, :]=True` and `data.mask[:, 1]=True` in Moments. These masks were removed when evaluating the model fit using Poisson residuals (see Section 5.2 below).

#### 5.1 Demographic model

We fit the observed joint SFS data to a simple demographic model of a population split with subsequent migration, implemented in the `moments.Demographics2D.split_mig` function in Moments. In this model, an ancestral population of constant size $N_a$ instantaneously splits into two populations $T_{\text{split}} \cdot \lambda$ generations ago, where $T_{\text{split}}$ is measured in years and $\lambda$ is the average growth rate from Eq. S2. The sizes of the new populations are $N_a \nu_1$ and $N_a \nu_2$, respectively, respectively, allowing for the total population size to increase or decrease after the split. We also allow for continuous migration between the two populations at a rate $m_0$ per individual per generation. This model constitutes a multi-population generalization of the two-epoch model employed by Mah *et al* [21] to study microbial demographic histories in several industrialized human cohorts. In particular, the addition of an explicit migration rate parameter allows us to quantify the timing and extent of genetic isolation between the Tsimane and Hadza microbiomes.

Since Moments outputs the inferred split time in population-scaled units ($\tilde{T}_{\text{split}} \equiv T_{\text{split}}\lambda/2N_a$), we applied the following steps to convert this estimate into years. We note that the ancestral population size ($N_a$) does not affect the relative probability density of the predicted SFS; it only determines the number of sites that are polymorphic in a population, which is proportional to the total number of sites ($L$). Moments provides the scaling factor $\theta \equiv 4N_a\mu L$ using the function `Inference.optimal_sfs_scaling`, which compares the model with the observed number of SNVs in the data. Since these SNVs are derived from the subset of sites that passed projection, we calculated $L$ as the number of synonymous sites sufficiently covered in the MAGs to meet the projection threshold. We then used the following formula to convert the inferred split time into years:

$$T_{\text{split}} = \frac{\tilde{T}_{\text{split}} \cdot 2N_a}{\lambda} = \tilde{T}_{\text{split}} \cdot \frac{\theta}{4\mu\lambda L} \cdot 2\,. \tag{S13}$$

using the yearly mutation rate of $\mu\lambda \sim 10^{-6.5}$ mutations per site per year estimated in Section 1.2.

#### 5.2 Quality control filters of model fits

In this section, we describe the quality control filters applied to refine the dataset for demographic inference, some of which are based on the results of model fits. These filters reduced the total number of species analyzed to the 15 that are shown in Fig. 4 in the main text. The number of species retained at each filtering step is summarized in a flowchart (Fig. S8). The inferred model parameters and estimated population split times are reported in Supplementary Table 4.

**Impact of clade structure.** Some bacterial species harbor complex population structures, such as distinct clades or subspecies, where horizontal gene transfer between clades occurs at lower rates. This complexity violates the assumptions of our simple demographic model and could bias the inferred split times. To focus on species with sufficiently simple population structures that align with the assumptions of our model, we applied a heuristic approach to detect and filter out species with strong clade structure. We applied hierarchical clustering to the pairwise distance matrix based on ANI differences, using the `scipy.hierarchy.fcluster` function with the criterion set to `'maxclust_monocrit'` to form two maximally separated clusters. To be conservative, we excluded a species from the Moments analysis if the smaller of the two clusters contained more than one MAG.

To illustrate the impact of clade structure on model fit and inference, we compared both the mean residual sizes and the inferred split times between species identified as clade-structured and those without clade structure (Fig. S9). Here, residuals refer to the difference between the model prediction and the observed SFS, calculated using `moments.Inference.linear_Poisson_residual`. The Poisson residual is defined as $(\mathrm{model} - \mathrm{data})/\sqrt{\mathrm{model}}$, where "model" and "data" represent individual elements of the predicted and observed SFS.

We found that species without clade structure tend to have smaller residuals, indicating better model fit, while clade-structured species show much larger inferred split times as expected. This discrepancy is likely due to the larger and older genetic diversity maintained by the presence of two or more clades, which can inflate the inferred split time defined by geography alone. These results highlight the importance of controlling for population structure during demographic inference.

**Filters based on uncertainty and residual size** One internal check for model fit is the uncertainty in parameter estimates provided by Moments. Moments calculates this uncertainty using the inverse of the Godambe Information Matrix [22]. In practice, we observed that the uncertainty in some parameter estimates can exceed the magnitude of the parameter itself, potentially indicating model misspecification or poorly constrained parameters. To address this, we retained only species with well-defined uncertainties, specifically those where the ratio of uncertainty to inferred split time was smaller than 0.5.

In addition, we applied a filter based on the mean absolute residual size to focus on species with the best model fits. We computed the mean absolute residual size for the two-dimensional joint SFS and the single-population (marginalized) SFS. We used a threshold of mean absolute residual size $< 2$ to exclude species with poor fits. Figure S10 shows the mean absolute residual sizes for the 27 species that passed the clade structure and uncertainty filters above.

#### 5.3 Visualizing model fits

Moments predicts a two-dimensional joint SFS (Figs. S11 and S12), but the high dimensionality of this distribution makes it challenging to visualize differences between the data and the inferred model, as well as between different demographic scenarios. To assess model fit, we first examined traditional diagnostics, including the marginalized SFS for each population, direct comparisons of observed and predicted joint SFS heatmaps, and the distribution of residuals across observed SNV frequencies.

These diagnostics, which are shown in Figs. S11 and S12, demonstrate overall agreement between model predictions and observed data. However, they do not directly reveal signals of population isolation, making it hard to directly distinguish the null scenario of no population split (free migration) from the inferred model and the data.

To address this limitation, we also employed a summary statistic for the two-dimensional SFS based on the frequency of "population-specific SNVs" – variants that are exclusively observed in only one of the two populations. The rationale is that population isolation reduces the chances that a recently produced SNV will be shared between the two populations. Importantly, the probability of this event will depend on the total frequency of the SNV across populations, making this statistic particularly sensitive to the effects of isolation.

The probability that a SNV is observed in only one population depends on three main factors. First, SNVs at high frequencies tend to have persisted in the population for longer, increasing the likelihood of migration between populations. Second, high-frequency SNVs are more likely to have been present in the ancestral population prior to the split, making them more likely to appear in both populations. Lastly, low-frequency SNVs are less likely to be sampled and may appear exclusive to one population by chance. Together, these factors contribute to a characteristic decay in the fraction of population-specific SNVs as total population frequency increases.

We tested this idea by plotting the fraction of population-specific SNVs as a function of total population frequency, using simulated SFS data for the same demographic model applied in our Moments inference. Figure S13 illustrates how this relationship varies with population split time and migration rate between populations. As expected, the sharpest decay in population-specific SNVs was observed in the no-separation scenario ($\tilde{T}_{\text{split}} = 0$). In simulations without migration following the split (population-scaled migration rate $m \equiv 2N_a m_0 = 0$), the decay was highly sensitive to split time, effectively distinguishing different separation times below $\tilde{T}_{\text{split}} = 2$. However, with moderate migration ($m = 1$), the curve still distinguished shorter split times but lost sensitivity for splits older than $\tilde{T}_{\text{split}} = 1$. At higher migration rates ($m = 5$), the ability to distinguish split times diminished further, with the curves approaching the no-split (or "free migration") limit.

Having demonstrated the utility of this statistic with simulations, we next applied it to the 15 species with the best Moments fits. In the majority of species, this statistic revealed a clear distinction between the observed data and the no-split scenario (Figs. 4B and S14), highlighting the robust effect of the population split. Additionally, we observed strong agreement between the predicted curves from the inferred model and the data, validating the model's ability to capture key patterns in the site frequency spectrum.

Interestingly, for the handful of species that were also analyzed by Mah *et al* [21] — *Ruminococcus bromii*, *Ruminococcus instestinalis*, and *Faecalibacterium prausnitzii* — we found that the Tsimane-Hadza split times inferred by Moments approximately coincided with the population expansions inferred by Mah *et al* using strains from industrialized human cohorts. The reasons for this correspondence remain unclear, but would be interesting to explore in future analyses combining industrialized and non-industrialized populations.

[18] Pauli Virtanen, Ralf Gommers, Travis E. Oliphant, Matt Haberland, Tyler Reddy, David Cournapeau, Evgeni Burovski, Pearu Peterson, Warren Weckesser, Jonathan Bright, Stéfan J. van der Walt, Matthew Brett, Joshua Wilson, K. Jarrod Millman, Nikolay Mayorov, Andrew R. J. Nelson, Eric Jones, Robert Kern, Eric Larson, C J Carey, İlhan Polat, Yu Feng, Eric W. Moore, Jake VanderPlas, Denis Laxalde, Josef Perktold, Robert Cimrman, Ian Henriksen, E. A. Quintero, Charles R. Harris, Anne M. Archibald, Antônio H. Ribeiro, Fabian Pedregosa, Paul van Mulbregt, and SciPy 1.0 Contributors. SciPy 1.0: Fundamental Algorithms for Scientific Computing in Python. *Nature Methods*, 17:261–272, 2020.

[19] Julien Jouganous, Will Long, Aaron P Ragsdale, and Simon Gravel. Inferring the Joint Demographic History of Multiple Populations: Beyond the Diffusion Approximation. *Genetics*, 206(3):1549–1567, July 2017.

[20] Santiago G. Medina-Muñoz, Diego Ortega-Del Vecchyo, Luis Pablo Cruz-Hervert, Leticia Ferreyra-Reyes, Lourdes García-García, Andrés Moreno-Estrada, and Aaron P. Ragsdale. Demographic modeling of admixed Latin American populations from whole genomes. *The American Journal of Human Genetics*, 110(10):1804–1816, October 2023.

[21] Jonathan C Mah, Kirk E Lohmueller, and Nandita R Garud. Inference of the Demographic Histories and Selective Effects of Human Gut Commensal Microbiota Over the Course of Human History. *Molecular Biology and Evolution*, 42(2):msaf010, February 2025.

[22] Alec J. Coffman, Ping Hsun Hsieh, Simon Gravel, and Ryan N. Gutenkunst. Computationally Efficient Composite Likelihood Statistics for Demographic Inference. *Molecular Biology and Evolution*, 33(2):591–593, February 2016.

#### Supplementary Figures

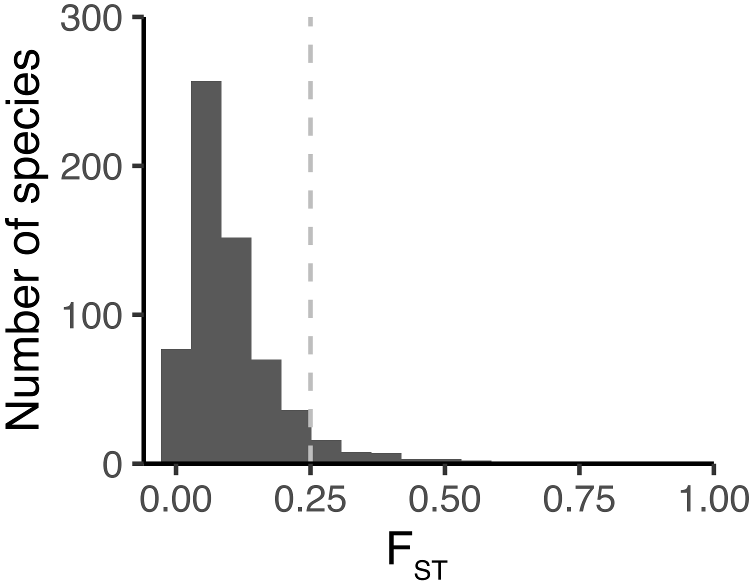

Figure S1: Distribution of $F_{ST}$ values between microbial strains sampled from Tsimane and Hadza individuals, calculated using Eq. S7. For comparison, the dashed line indicates the approximate $F_{ST}$ between African and Native American human populations.

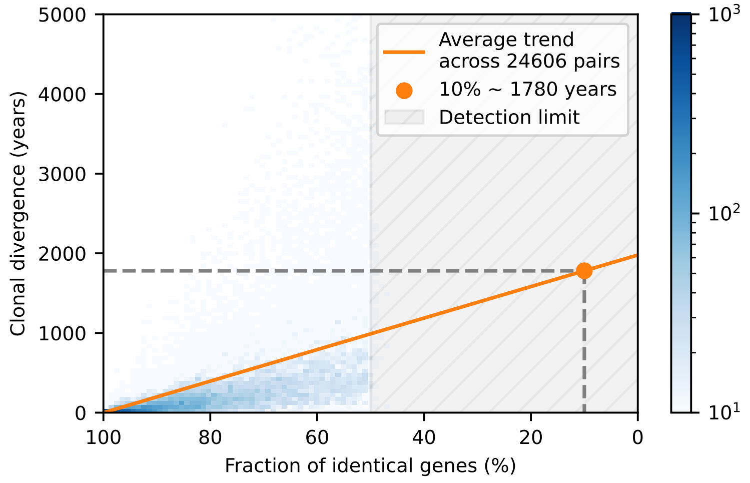

Figure S2: 2-D histogram (density heatmap) showing the linear trend between the inferred clonal divergence time and the fraction of identical genes. Solid line shows the linear fit using the data of all 57,505 MAG pairs with more than 50% identical genes, aggregated across species. Symbol denotes the linearly extrapolated divergence time corresponding to 10% identical genes. Color bar indicates the number of MAG pairs within each bin.

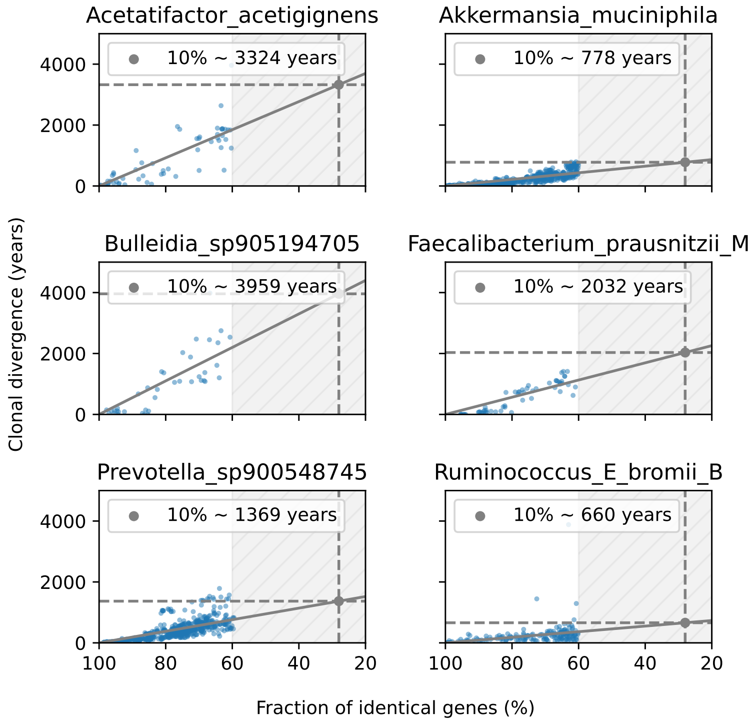

Figure S3: Scatterplots analogous to Fig. S2, showing the linear trend for six example species. Each blue dot denotes one pair of MAGs.

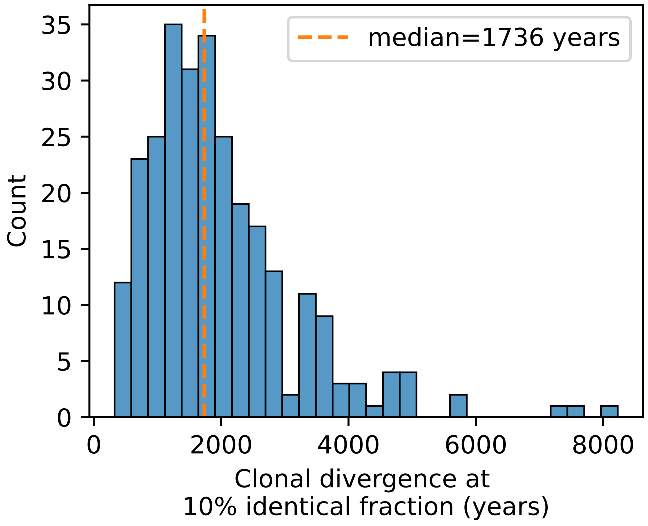

Figure S4: Histogram of the extrapolated clonal divergence times corresponding to 10% identical genes, for each microbial species.

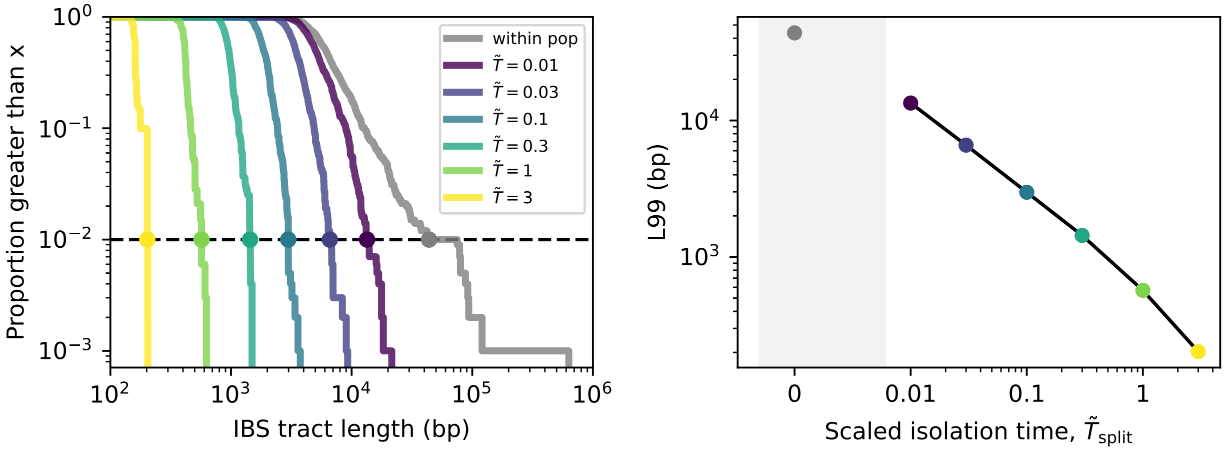

Figure S5: Left: IBS tract length distributions of simulated microbial genomes using FastSimBac. Curves show the reverse cumulative distributions of the longest IBS tract aggregated over 1000 pairs of sequences. All distributions are computed for between-population genome pairs,except for the "within pop" curve. Dashed line and circles indicate the tract lengths at the 99th percentile (L99). Right: Same data as the left panel, showing the L99 lengths as a function of the scaled isolation time between the populations in the simulation. Symbol at zero isolation time indicates the value observed for within-population pairs.

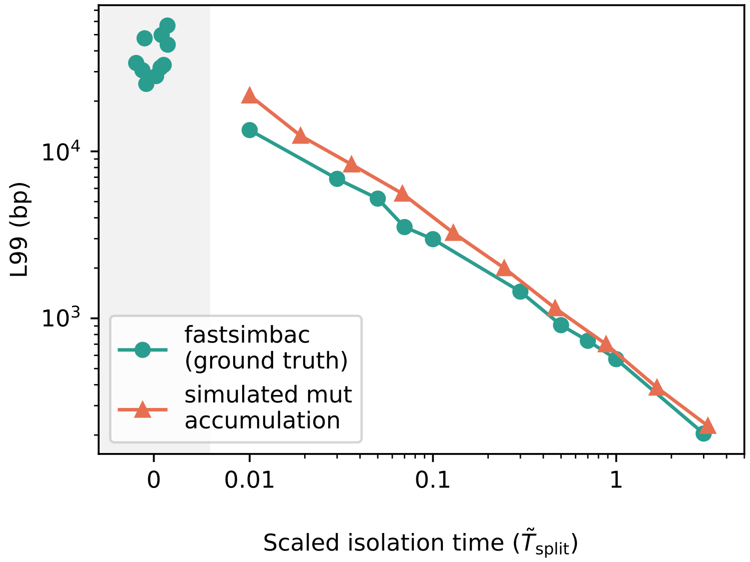

Figure S6: Comparison between the ground truth L99-$T_{\text{split}}$ relationship observed in FastSimBac simulations, and the approximate version obtained from the mutation accumulation approach in Section 4.3. The x-axis represents the scaled isolation time, and the y-axis shows L99 length. The ground truth, derived from full recombination-based FastSimBac simulations, is shown in cyan, while the approximate mutation accumulation simulation is shown in orange. The close agreement between the two curves suggests that mutation accumulation alone provides a reasonable first-order approximation of IBS tract dynamics over time, particularly at longer isolation times.

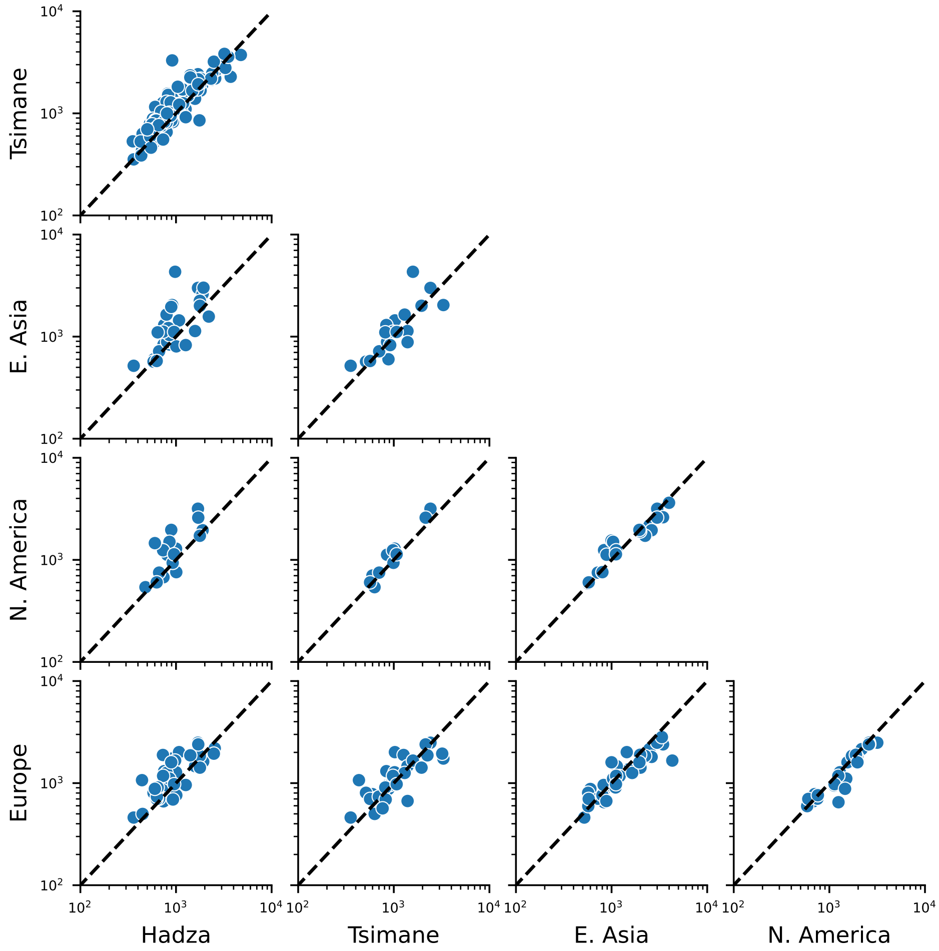

Figure S7: Scatterplots comparing the observed within-population L99 lengths between different pairs of populations. Each point indicates a single species. Dashed lines indicate the equality line ($y = x$). For each panel, only species with more than 200 within-population pairs in both populations are included.

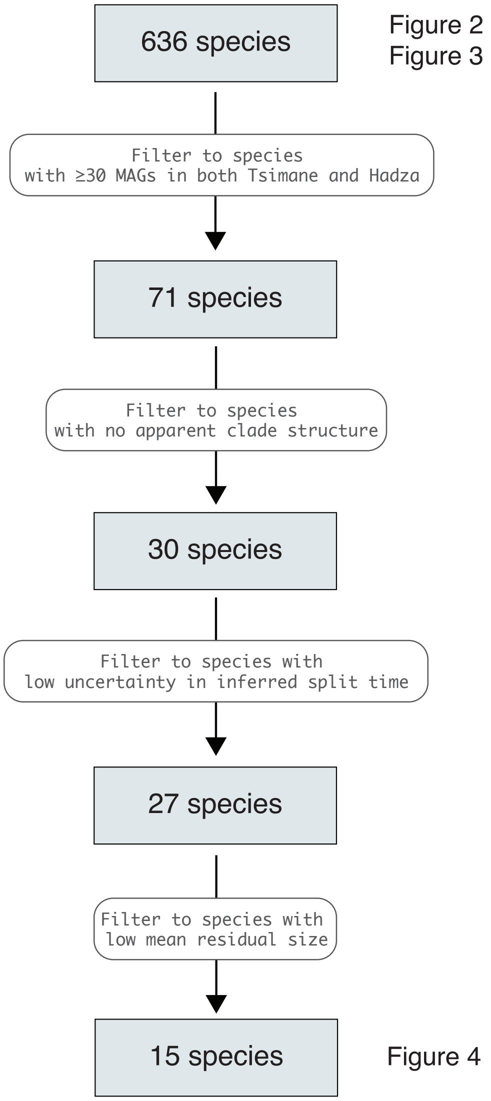

Figure S8: Flowchart summarizing the data filtering steps used for the Moments analysis (Section 5.2).

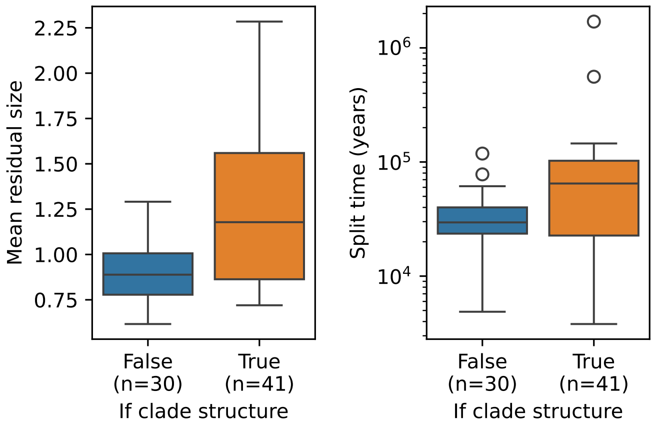

Figure S9: Box plots comparing the mean residual sizes (left) and inferred split times (right) of demographic model fits for species with and without clade structure, as identified by our clade detection method. The sample size $n$ denotes the number of species in each bin (note: these species have not yet been filtered for parameter uncertainty or residual size, as described in Section 5.2). These data show that species without clade structure exhibit smaller residuals, indicating better alignment with the assumptions of the simple demographic model in Section 5.1. Species with clade structure also show higher inferred split times, potentially reflecting increased genetic diversity within species.

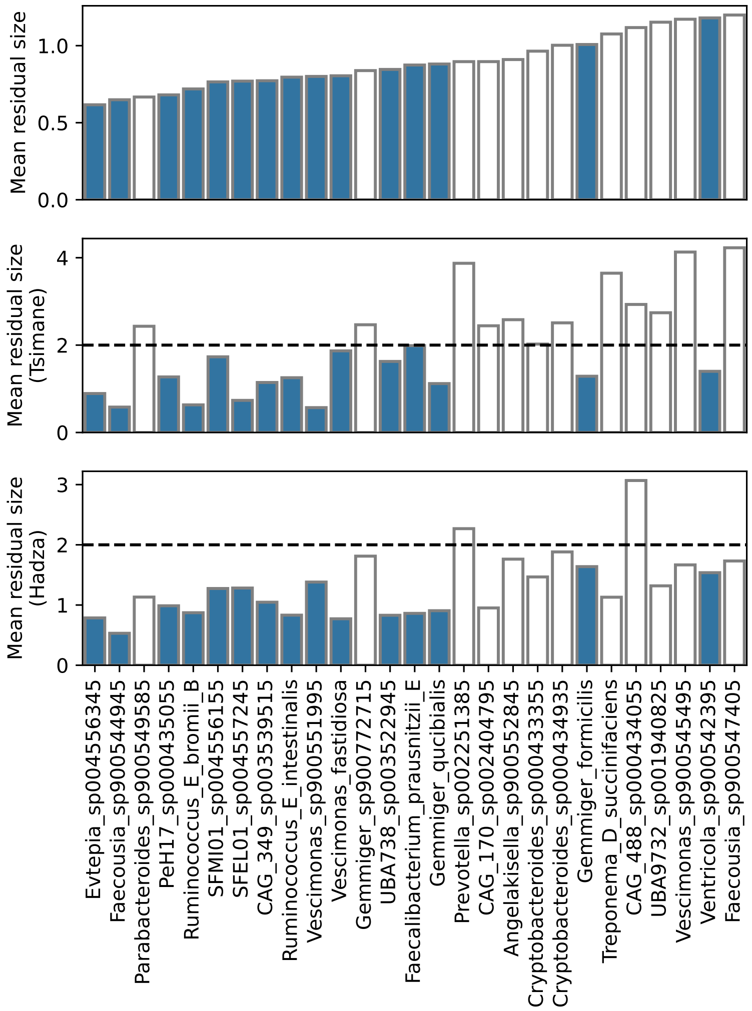

Figure S10: Bar plot showing the mean absolute residual sizes for the 27 species that passed the clade structure and uncertainty filters. The top panel shows the mean residual for the joint SFS, while the bottom two panels show the mean residual of the single-population (marginalized) site frequency spectrum. The dashed line represents the residual size filter threshold of 2. Solid bars denote species that passed the residual size filter.

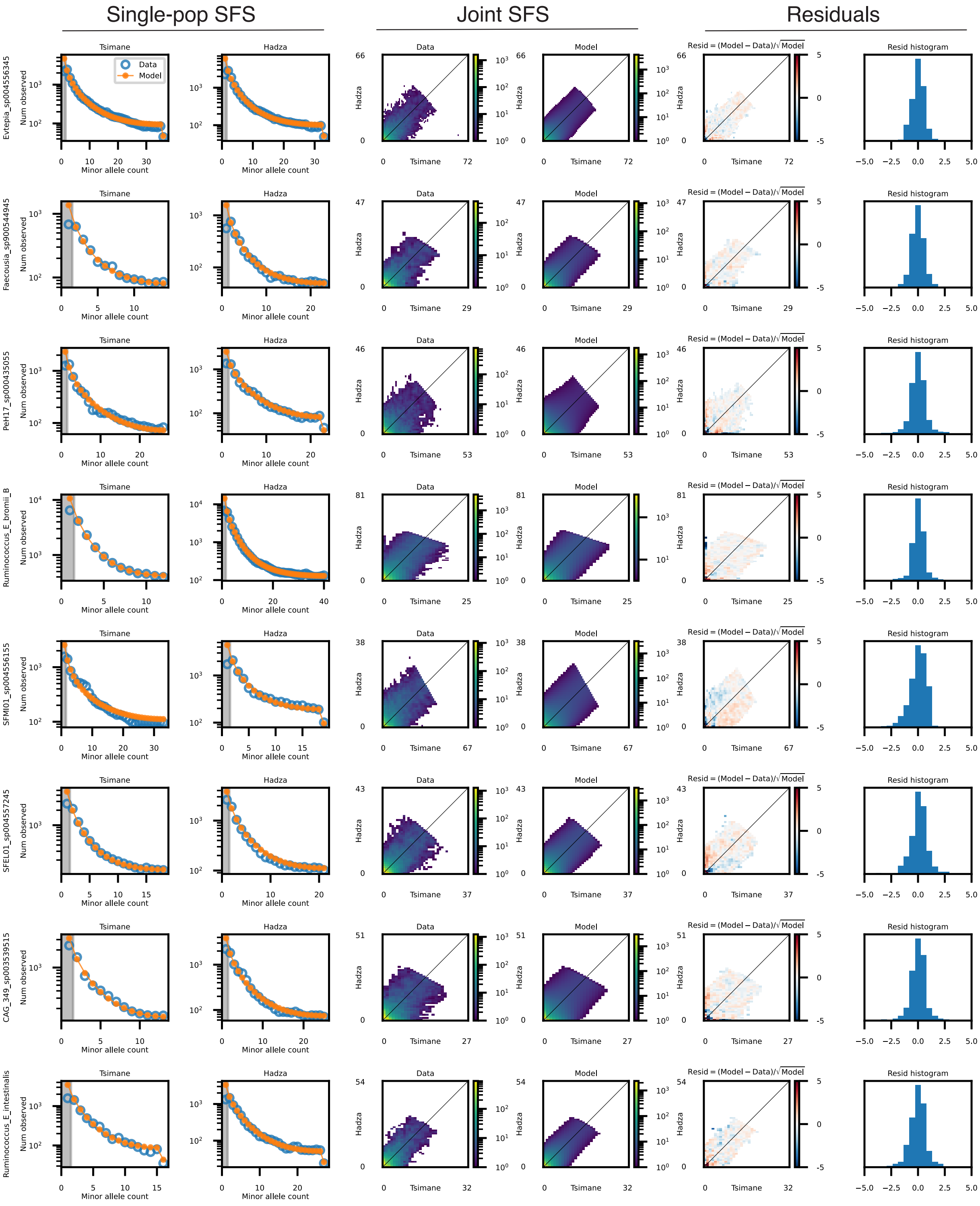

Single-pop SFS
Joint SFS
Residuals
Tsimane
Hadza
Data
Model
Resid = (Model − Data)/√Model
Resid histogram
Num observed
Minor allele count
Evtepia_sp004556345
Faecousia_sp900544945
PeH17_sp000435055
Ruminococcus_E_bromii_B
SFMI01_sp004556155
SFEL01_sp004557245
CAG_349_sp003539515
Ruminococcus_E_intestinalis

Figure S11: Summaries of demographic model fits inferred by Moments for 8 of the 15analyzed species. The remaining species are shown in Fig. S12. Each row represents one species. The first two columns show the single-population (marginalized) site frequency spectrum (SFS) for the Tsimane and Hadza microbial populations, respectively. The shaded region indicates singleton SNVs (minor allele count = 1), which are masked during model fitting. Columns 3 and 4 show heatmaps of the joint SFS between Tsimane and Hadza microbial populations for observed data and model predictions. Column 5 shows heatmap of residuals between the model predictions and observed data. Column 6 shows a histogram of the residual values from column 5, with the y-axis representing probability density on a linear scale (ticks omitted for compactness).

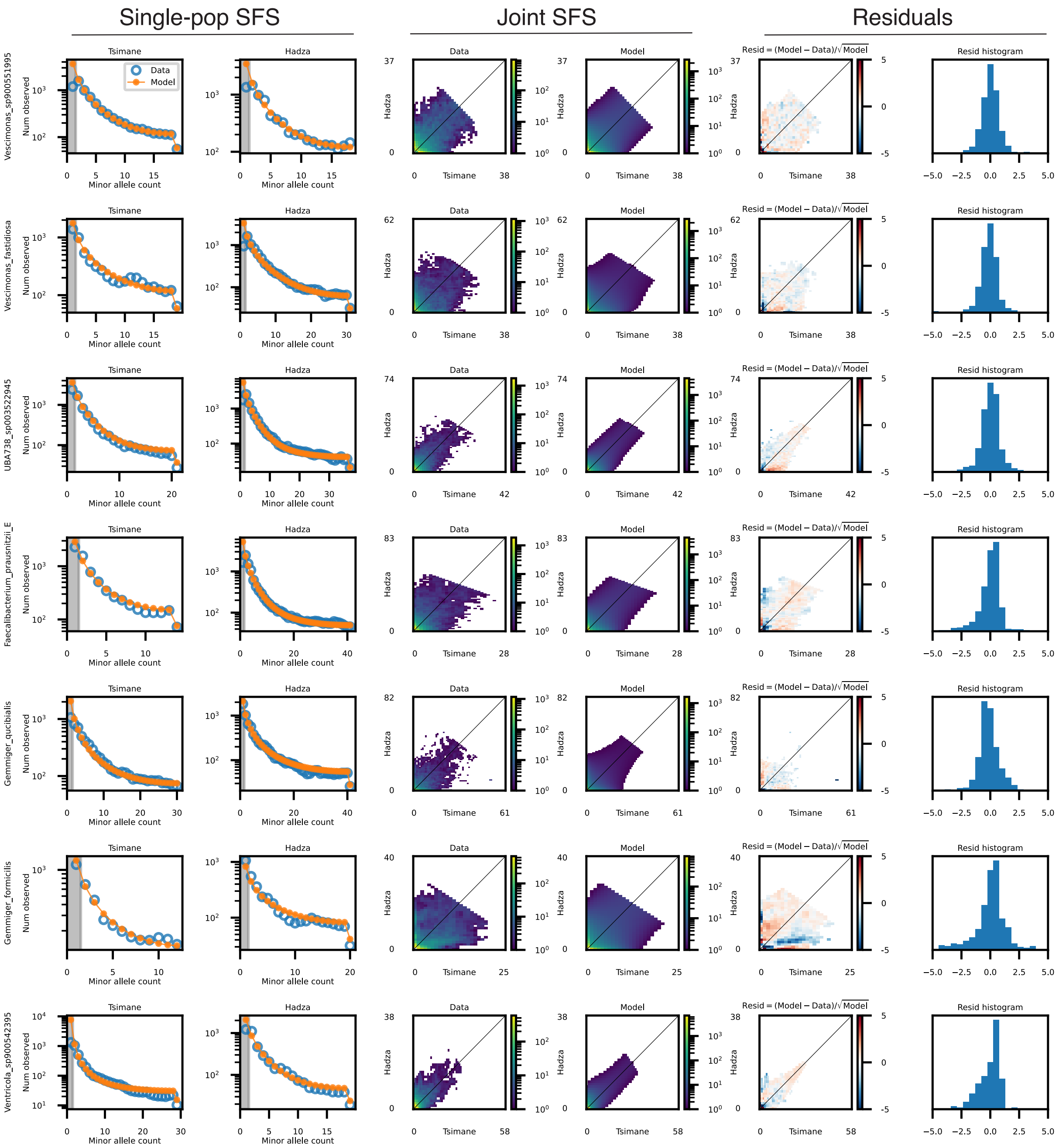

Figure S12: Same as in Fig. S11, but for the remaining seven species.

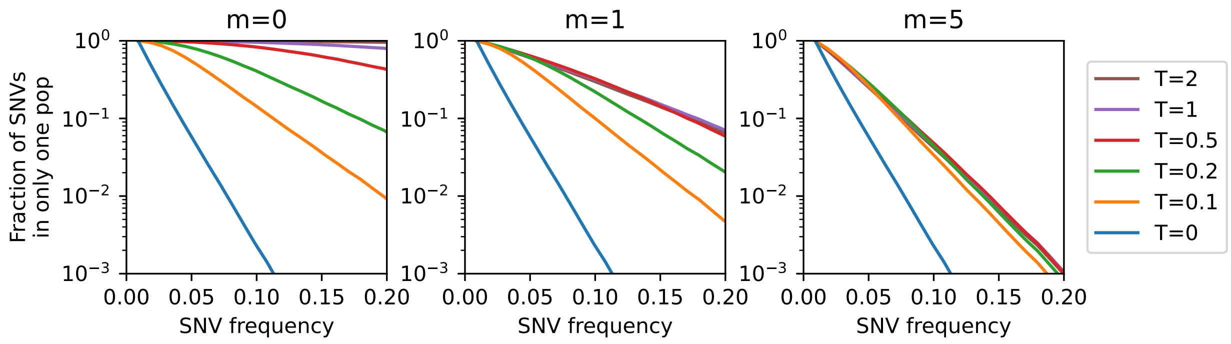

Figure S13: Plots showing the predicted fraction of population-specific SNVs for simulated data, analogous to Figure 4B in the main text. Curves are computed using the SFS simulated by the `split_mig` demographic model, with parameters $\nu_1 = \nu_2 = 2$ and varying split times and migration rates. Sample sizes were chosen to match those of the species *UBA738 sp003522945*. The parameter $m$ represents the scaled migration rate, defined as $2N_a m_0$.

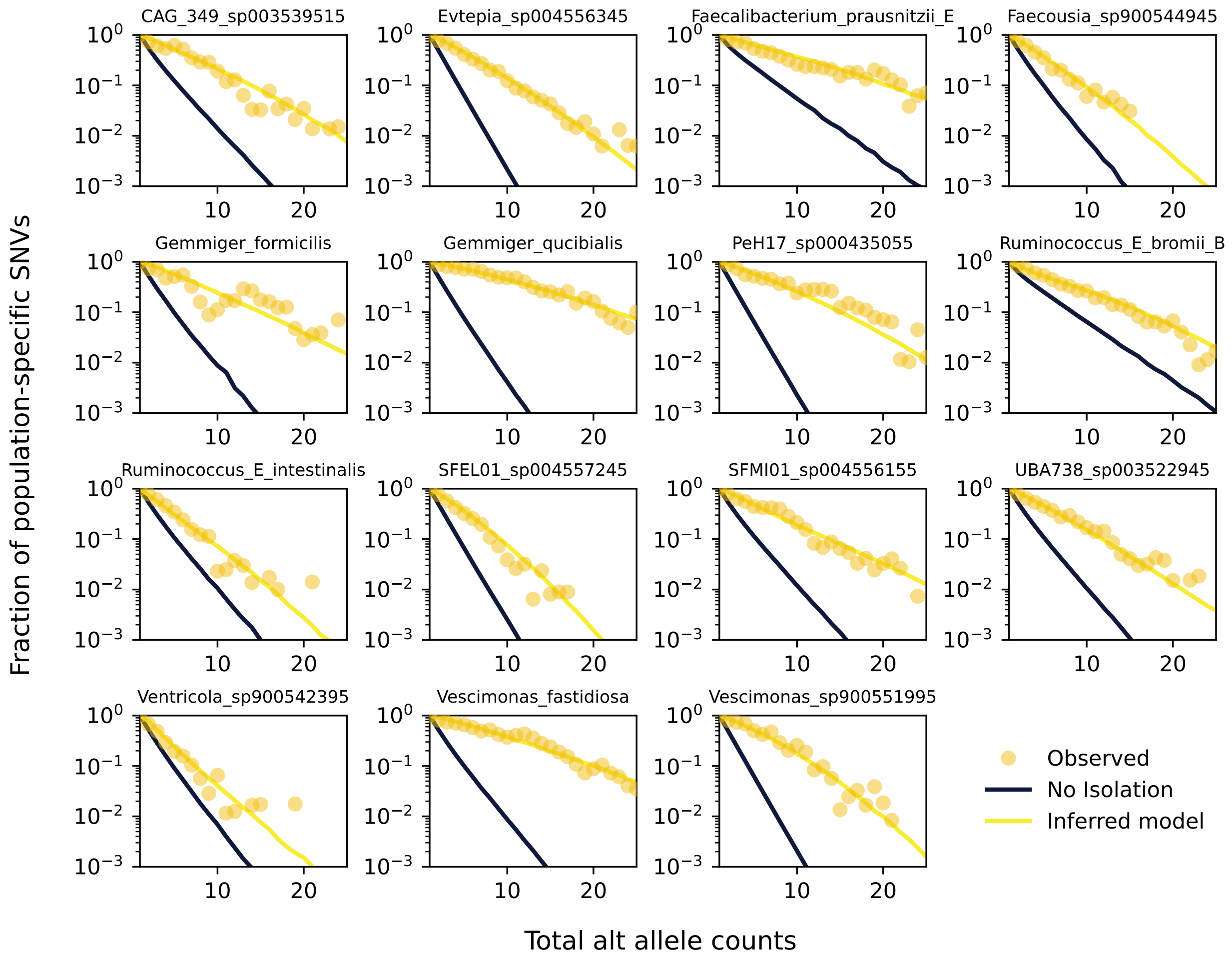

Figure S14: Analogous to Figure 4B in the main text. The yellow curve represents the prediction from the demographic model inferred using the full SFS, while the black curve corresponds to the null scenario of no population split. The null curve is computed by modeling the distribution of alternative alleles between populations using a binomial distribution, weighted by their relative population sizes. This is equivalent to randomly reshuffling alternative alleles between populations, simulating a scenario of free migration.
